# PfGCN5, a global regulator of stress responsive genes, modulates artemisinin resistance in *Plasmodium falciparum*

**DOI:** 10.1101/679100

**Authors:** Mukul Rawat, Abhishek Kanyal, Aishwarya Sahasrabudhe, Shruthi S. Vembar, Jose-Juan Lopez-Rubio, Krishanpal Karmodiya

## Abstract

*Plasmodium falciparum* has evolved resistance to almost all front-line drugs including artemisinins, which threatens malaria control and elimination strategies. Oxidative stress and protein damage responses have emerged as key players in the generation of artemisinin resistance. In this study, we show that PfGCN5, a histone acetyltransferase, binds to the stress responsive and multi-variant family genes in poised state and regulates their expression under stress conditions. We have also provided biochemical and cellular evidences that PfGCN5 regulates stress responsive genes by acetylation of PfAlba3. Furthermore, we show that upon artemisinin exposure, genome-wide binding sites for PfGCN5 are increased and it is directly associated with the genes implicated in artemisinin resistance generation like BiP and TRiC chaperone. Moreover, inhibition of PfGCN5 in artemisinin resistant parasites, Kelch13 mutant, K13I543T and K13C580Y (RSA∼ 25% and 6%, respectively) reverses the sensitivity of the parasites to artemisinin treatment indicating its role in drug resistance generation. Together, these findings elucidate the role of PfGCN5 as a global chromatin regulator of stress-responses with potential role in modulating artemisinin drug resistance, and identify PfGCN5 as an important target against artemisinin resistant parasites.

**Author Summary:** Malaria parasites are constantly adapting to the drugs we used to eliminate them. Thus, when we use the drugs to kill parasites; with time, we select the parasites with the favourable genetic changes. Parasites develop various strategies to overcome exposure to the drugs by exhibiting the stress responses. The changes specific to the drug adapted parasites can be used to understand the mechanism of drug resistance generation. In this study, we have identified PfGCN5 as a global transcriptional regulator of stress responses in *Plasmodium falciparum*. Inhibition of PfGCN5 reverses the sensitivity of the parasites to the artemisinin drug and identify PfGCN5 as an important target against artemisinin resistant parasites.

## Introduction

Malaria is a life threatening infectious disease caused by parasites from the genus *Plasmodium,* with an estimated 200 million cases worldwide [1]. The *Anopheles* mosquito serves as a vector for varied species of the human malaria parasite namely *P. falciparum, P. vivax, P. ovale, P. malariae* and *P. knowlesi*. Of these five species, *P. falciparum* causes most lethal form of malaria. The *Plasmodium* life cycle consists of two phases, sexual and asexual in mosquitoes and humans, respectively. Since *Plasmodium* completes its life cycle in two different hosts, it requires mechanisms for coordinated modulation of gene expression [2]. An efficient transcriptional and post-transcriptional regulation of gene expression enables it to establish chronic infection in humans [3, 4]. Moreover, recent studies have attested the importance of epigenetic mechanisms in regulation of gene expression [3, 5, 6]. Morphological changes observed during the development of the malaria parasite in erythrocytes are also governed by the fine-tuning of gene expression [2]. Furthermore, most of the genes in *Plasmodium* are reported to be poised (genes exhibit high levels of histone activation marks but no transcription), which favors the plasticity of its gene expression programs [5, 7, 8].

During the asexual life cycle, when *Plasmodium* is developing within mature red blood cells (RBCs), it is exposed to different kinds of environmental and physiological stresses. For instance, during the trophozoite stage (18 h post-erythrocyte invasion (hpi)), the parasite converts hemoglobin to hemozoin, leading to the accumulation of reactive oxygen species (ROS) and consequently oxidative stress [9]. Various drugs like artemisinin, arteether etc. used for antimalarial treatment also lead to similar ROS build up, ultimately killing the parasite [9–11]. Another characteristic of malarial infection is the acute cyclical episodes of fever, with an increase in temperature to 41 degrees Celsius for about 2-6 hours. This periodic febrile response is triggered by the release of merozoites from RBCs [12, 13]. Since *Plasmodium* faces these stress conditions during each of its infectious cycle, it has possibly evolved mechanisms to resist the metabolic perturbations caused thereby. Interestingly, these stress responses are also known to mediate resistance against various antimalarial drugs [14, 15]. Several classical antimalarial drugs like chloroquine [16], sulfadoxine-pyrimethamine [17] and mefloquine [18] are no longer effective against *P. falciparum* [19, 20]. Currently, artemisinin based combination therapy is considered as the last line of defense against *P. falciparum* malaria [20]. However, since 2009, alarming reports of resistance against artemisinin have emerged in Southeast Asia. This region has historically served as the epicenter for emergence of anti-malarial drug resistance [21–23]. Recent reports from Eastern India have also suggested the presence of artemisinin resistant parasites based on pharmacokinetic and genetic parameters like increased parasite clearance half-life and novel Kelch13 mutations [24, 25].

Currently, there is limited knowledge on the mechanism by which the parasites develop resistance against artemisinin. Artemisinin resistant parasites are characterized by slow growth and reduced drug susceptibility at the ring stage of asexual growth [15]. Artemisinin resistant parasites are also shown to have extensive transcriptional deregulation, with transcriptional regulators emerging as important players in the evolution of drug resistance [26–29]. Multiple transcriptomics studies have revealed dormancy, oxidative stress response and protein metabolism to be key players in mechanism of artemisinin drug resistance generation [22, 26, 27, 29–31]. Unfortunately, the global transcriptional regulators of drug resistance generation remain unexplored in *P. falciparum*. Previous studies in higher (e.g., humans) and lower (e.g., *Toxoplasma gondii*) eukaryotes demonstrated that GCN5, a histone acetyltransferase plays an important role during stress conditions, where it has been associated with high level of transcriptional reprogramming required for stress adaptation [32–35]. GCN5 is conserved in *Plasmodium* species and till date, only two subunits of the GCN5 complex, namely PfGCN5 and PfADA2 are identified [36, 37]. Previous studies using DNA microarray have suggested that there is a weak but positive correlation between PfGCN5 and H3K9ac mark [38]. In this study, we dissected role of PfGCN5 under various physiological stress conditions in *P. falciparum* during intraerythrocytic development cycles. With the help of chromatin immunoprecipitation coupled high-throughput sequencing (ChIP-seq) and transcriptomic (RNA-sequencing) analyses, we show that PfGCN5 activates genes that are important for the maintenance of parasite cellular homeostasis during various stress conditions. Furthermore, we elucidate the role of PfAlba3 as the mediator of PfGCN5-dependent regulation of stress responsive genes. Collectively, our data identify histone acetyltransferase, PfGCN5 as a key chromatin regulator of stress responsive genes and reveals its important role in emergence of artemisinin drug resistance.

## Results

### PfGCN5 is associated with virulence and stress responsive genes

While paralogs of GCN5 are well studied in multiple systems, little is known about the function of GCN5 in *P. falciparum*. PfGCN5, encoded by *PF3D7_0823300,* contains histone acetyltransferase (HAT) and bromo (for binding to acetylated histones) domains at its C-terminal end (S1A Fig) [37]. To gain further insight into the function of PfGCN5 during asexual growth, we generated polyclonal antibodies against recombinant C-terminal HAT and bromo domains (amino acid 1183-1448) of PfGCN5 (also called α-HAT antibody, S1A-S1B Fig). We also raised polyclonal antibodies against a peptide from the N-terminal region of PfGCN5 (amino acids 9-25; called α-peptide antibody; S1A Fig). Specificity of the generated antibodies was determined by Western blotting using parasite lysate (S1C Fig), and by immunoprecipitation coupled with mass spectroscopy.

Next, to comprehend the transcriptional regulation mechanisms of PfGCN5, we performed chromatin immunoprecipitation coupled high-throughput sequencing (ChIP-seq) using the α-HAT and α-peptide PfGCN5 antibodies. ChIP-seq was performed at early trophozoite stage (24 hpi) of parasite growth as PfGCN5 exhibits high mRNA expression at this stage (S2A Fig) [39]. Peaks of local enrichment of PfGCN5 were determined after sequence alignment and normalization to input sequences using the MACS2 peak calling software. In total, we identified 754 high confidence common binding sites (fold enrichment >= 2; q value < 0.1) with α-HAT and α-peptide PfGCN5 antibodies, which corresponds to 403 genes (S2B Fig). The PfGCN5 bound sites using α-peptide PfGCN5 antibody are mentioned in S1 Table. When we averaged PfGCN5 binding density for the α-HAT and α-peptide antibodies across the average gene structure of *P. falciparum*, we observed identical profiles for the two (S2C Fig). This confirmed that the antibodies are well-correlated and specifically recognize PfGCN5.

Since different gene sets in *P. falciparum* have distinct histone modification distribution profiles [5], we measured the signal density of H3K9ac, a histone modification that is known to be mediated by PfGCN5, and compared it to PfGCN5 density distribution (as measured by α-peptide antibody) across all 5712 *P. falciparum* genes. Interestingly, PfGCN5 was enriched at the 3’ end and centre of the gene body of the 403 target genes identified by MACS2 analysis. In contrast, these genes have the H3K9ac marks distributed along the entire gene body (Fig1A, top panel). When compared with the heterochromatin protein (PfHP1) occupancy, which uniformly coats chromosome ends that contain a majority of the multi-copy variant genes (var, rifin and stevor), we found that PfGCN5 exhibits specific binding to antigenic variation genes, as shown in the representative example for Chromosome 1 from the trophozoite stage (Fig 1B). These results corroborate our earlier findings where we have shown that stress and stimuli dependent genes show enrichment of histone modifications at the centre and towards the 3’-end of the genes [5], while genes belonging to other housekeeping functions demonstrate uniform distribution of histone modifications.

**Figure 1:**
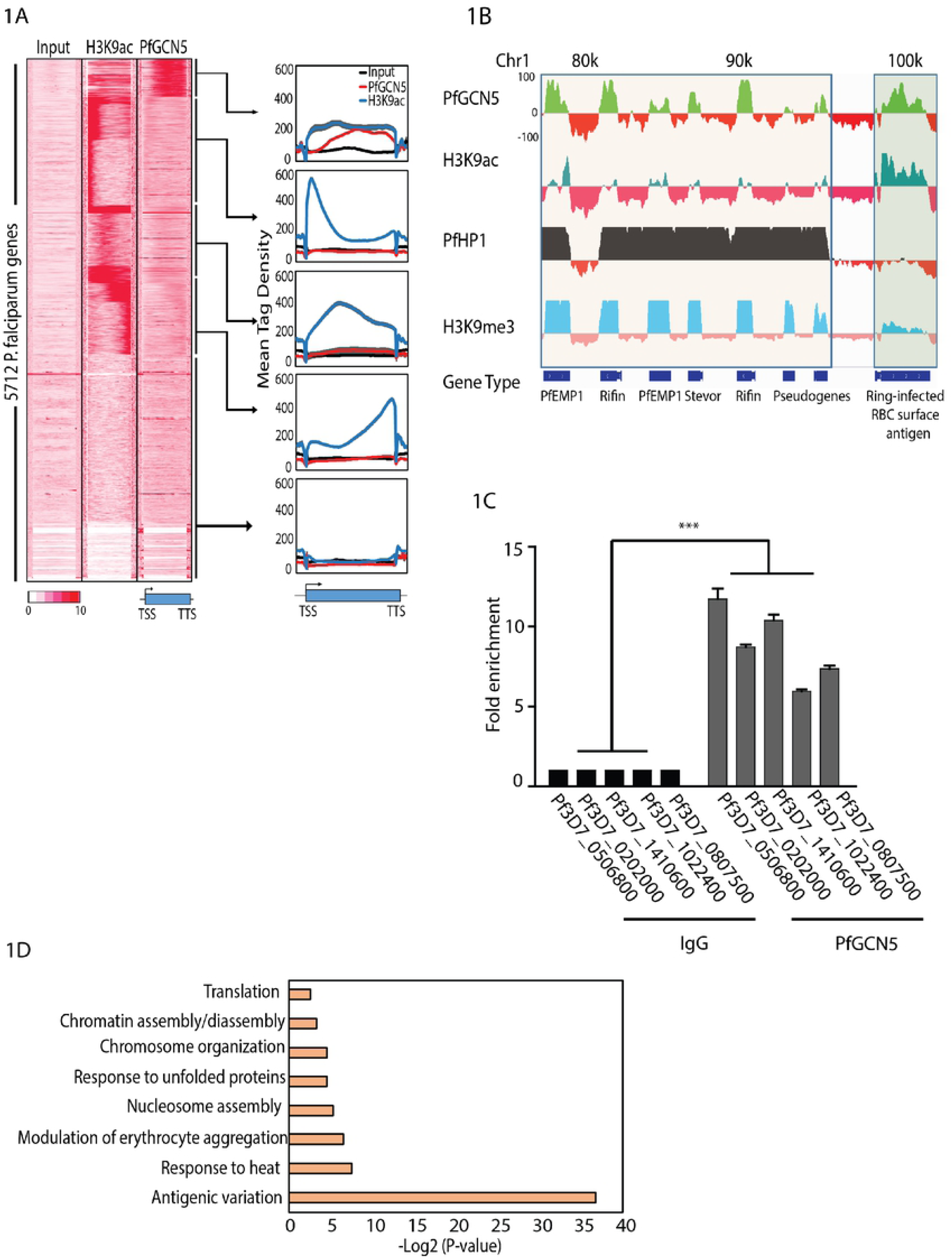
*Pf*GCN5 is associated with virulence and stress responsive genes. (A) Heat map showing the ChIP-seq tag counts at 5712 *P. falciparum* genes for H3K9ac and PfGCN5. PfGCN5 was enriched only over a subset of genes having H3K9ac enrichment indicating that it is not a general transcription co-activator. *Pf*GCN5 was found to be enriched mostly at the 3’ end of the genes and towards the centre of the genes. (B) IGV browser snapshot of representative genes having PfGCN5 binding. Binding of H3K9ac, Heterochromatin protein 1 (HP1) and H3K9me3 are also represented for comparison on the same genes. (C) ChIP-qPCR of selected genes confirms PfGCN5 binding to ChIP-seq targets. The results are shown as fold enrichment of ChIP performed with PfGCN5 α-peptide antibody versus non-immune IgG. (D) Gene ontology for the genes which were found to be bound by PfGCN5 using ChIP sequencing. Antigenic variation and other genes required during stress conditions are overrepresented in gene ontology.

Further, to validate the peaks obtained in ChIP-seq, we performed ChIP-qPCR on randomly selected genomic loci enriched for PfGCN5 and confirmed its binding (Fig 1C). Lastly, gene ontology (GO) analysis of PfGCN5 bound genes indicated enrichment of terms such as antigenic variation, stress response to heat, and response to unfolded proteins (Fig 1D), suggesting that PfGCN5 may play a role in the regulation of stress responsive and stimuli-dependent genes in *P. falciparum*. The presence of an expanded repertoire of GCN5-related N-acetyltransferase (GNAT) family of histone acetyltransferases [40] clearly indicates the possibility of uncharacterized HATs as writers of H3K9ac mark in *P. falciparum*.

### PfGCN5 is not a general transcription co-activator; it is specifically associated with stress/stimuli associated genes

Next, we investigated how PfGCN5 binding relates to transcriptional activity of a gene at the trophozoite stage. We systematically calculated the enrichment levels of PfGCN5 and H3K9ac, a general activation mark, at the gene body of all *P. falciparum* genes and compared it to the relative expression levels of genes as evaluated by RNA-seq-based transcriptomic analysis. As expected, we observed a positive correlation between H3K9ac enrichment and the expression status of the downstream gene (Fig 2A; left panel). On the other hand, we did not observe strong positive correlation between PfGCN5 gene-body occupancy and the expression of nearby genes (Fig 2A; right panel). Genes with either high or low gene expression levels (outlier points for log2 read density) showed high PfGCN5 occupancy (Fig 2A), suggesting that PfGCN5 binds to both active and suppressed/poised genes. In order to confirm this, we compared the expression levels of genes bound by PfGCN5 and contrasted them with the expression of all the *P. falciparum* genes. The expression level of PfGCN5 bound genes spreads from high expression to low expression values (Fig 2B) indicating its presence on expressed as well as suppressed genes. Interestingly, many of the PfGCN5 bound genes have both activation (H3K9ac) and repression (H3K9me3) marks (Fig 2C), indicating suppressed yet poised for future activation. Thus, absence of global correlation with transcription and occupancy on suppressed/poised as well as active genes, suggest that PfGCN5 is not a general transcriptional co-activator rather it may specifically regulate stress responsive genes in *P. falciparum*.

**Figure 2:**
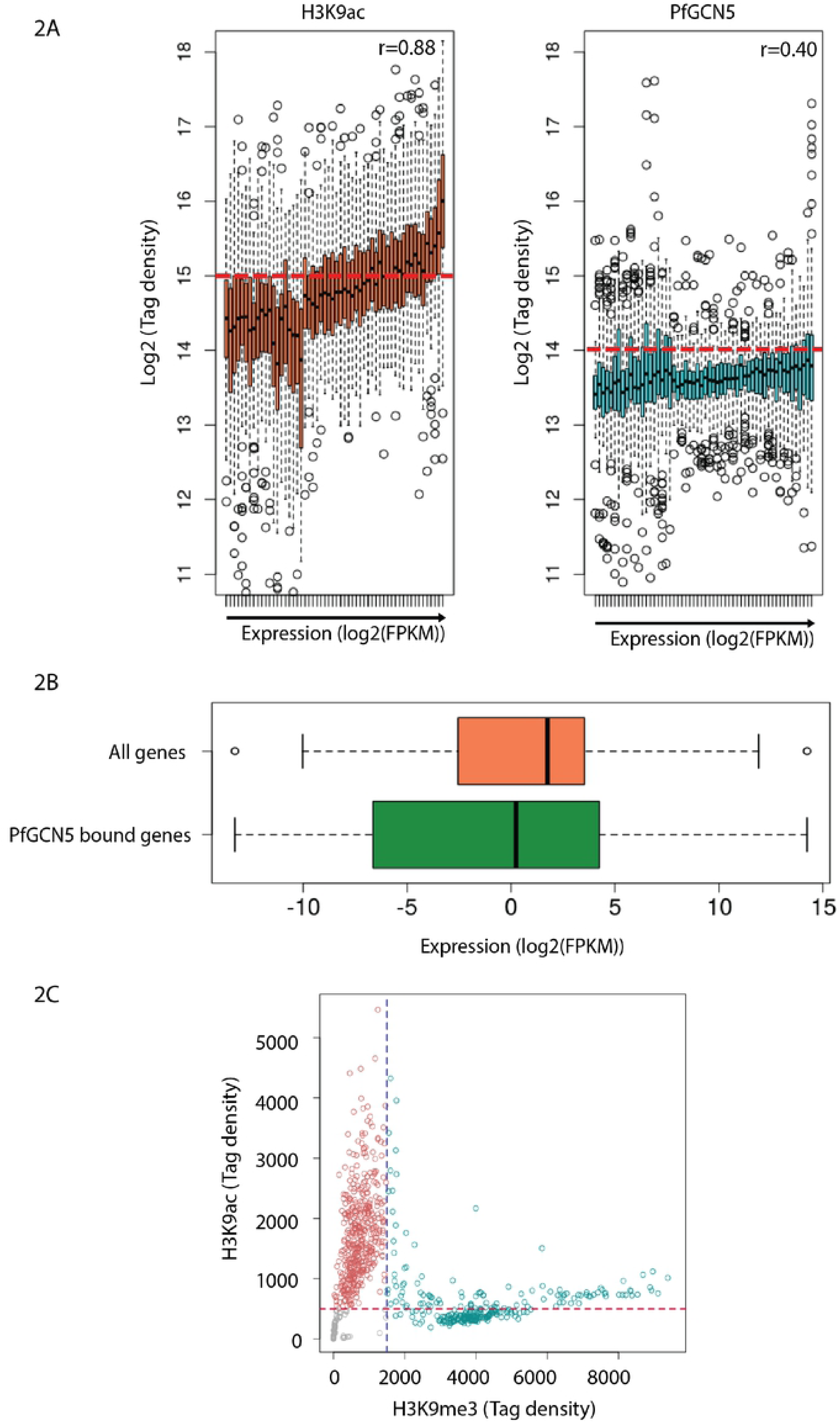
*Pf*GCN5 is not a general transcription coactivator; it is specifically associated with stimuli associated genes. (A) Box and whisker plots representing the correlation of genome-wide H3K9ac prevalence and PfGCN5 occupancy with the global gene expression. Absence of global correlation was found for the recruitment of *Pf*GCN5 and gene expression. This indicates that PfGCN5 is not a general transcription coactivator. (B) The expression level of the genes bound *Pf*GCN5 in comparison to all genes in *P. falciparum* is represented by the box plots. PfGCN5 is associated with highly as well as least expressed genes. (C) Scatter plot representing the correlation between the H3K9ac and H3K9me3 marks on PfGCN5 bound genes. The plot indicates that many genes having PfGCN5 binding also have H3K9me3 marks suggesting that these genes are either supressed or poised for future activation.

### PfGCN5 is a specific regulator of stress responsive genes

Next, we decided to look into the role of PfGCN5 during different stress conditions. Synchronized ring stage parasites were exposed to two different physiological stress conditions; temperature (40°C) and drug (30 nM artemisinin) exposure for 6 h. In order to confirm the stress response we looked at the expression level of marker genes, which are known to be upregulated during stress conditions in *Plasmodium*. For temperature stress we looked at the expression level of the heat shock protein, HSP70 (S3A Fig) [13, 41]. Since the artemisinin is known to induce oxidative stress through production of reactive oxygen species, we confirmed the oxidative stress by validating the expression levels of glutathione S-transferase and superoxide dismutase (S3A Fig) [42]. Interestingly, PfGCN5 was several fold upregulated upon physiological stress conditions as shown by quantitative real-time PCR (qPCR) using gene specific primers (Fig 3A). To identify the genes that are deregulated under these stress conditions, we performed transcriptomic analysis using RNA-sequencing and identified 727 and 942 genes (>2 fold change) deregulated upon artemisinin and high temperature exposure, respectively (Fig 3B-3C). Genes showing deregulation during stress conditions were also validated by qRT-PCR (S3B Fig). Most of the genes that are upregulated during both artemisinin and temperature stress conditions are reported to maintain cellular homeostasis (Fig 3B-3C; S2 Table). To further dissect the functional correlation between transcriptome deregulation and recruitment of PfGCN5 under different stress conditions, we performed ChIP-sequencing for PfGCN5 using α-peptide antibody during both temperature and artemisinin stress conditions. Notably, most of these genes which are bound by PfGCN5, are upregulated under artemisinin and temperature stress conditions (Fig 3D), indicating that PfGCN5 is associated with the activation of stress responsive genes.

**Figure 3:**
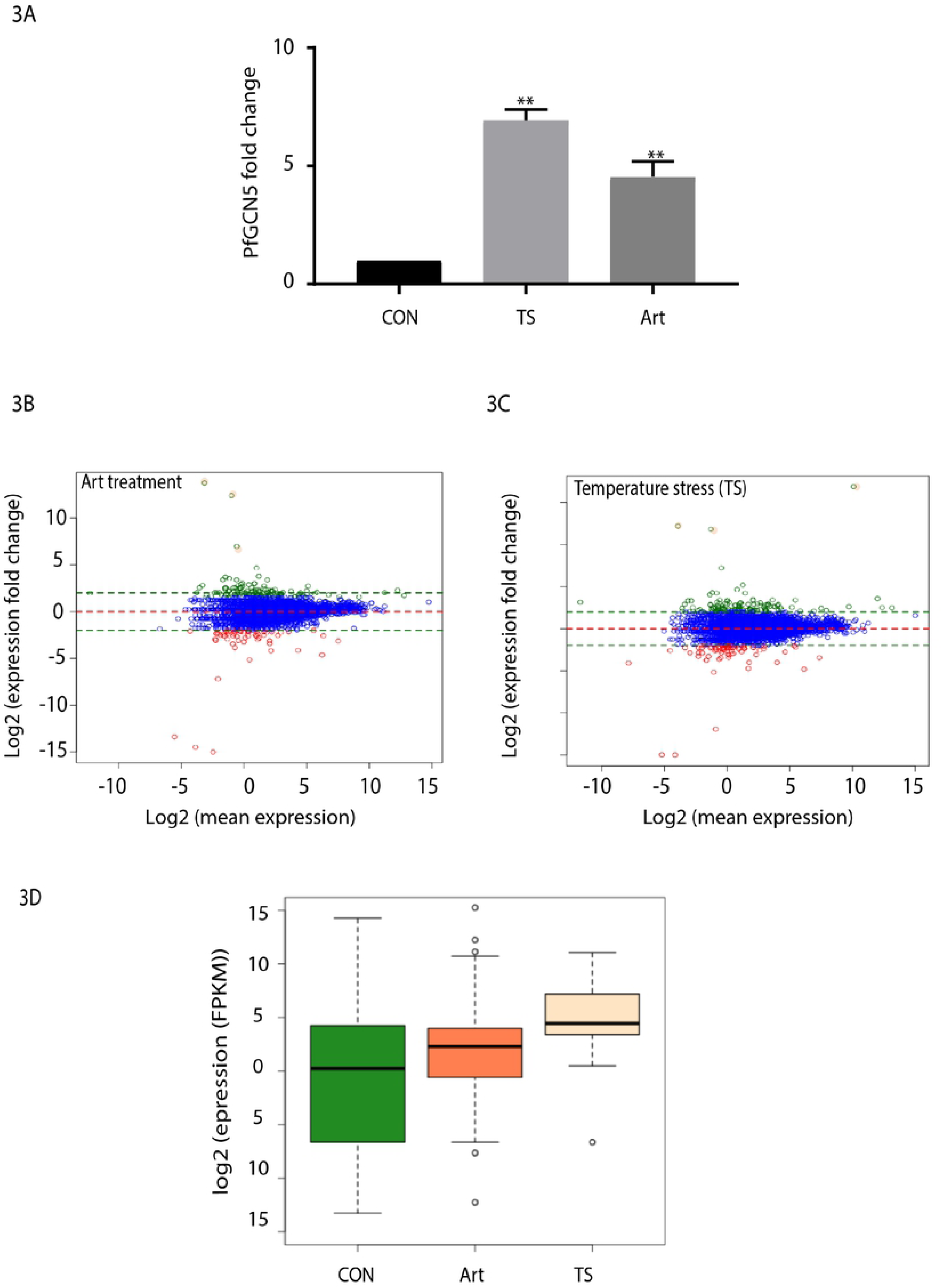
PfGCN5 is a specific stress regulator. (A) Change in the expression level of PfGCN5 during various stress conditions (N=3). PfGCN5 is found to be upregulated during heat stress and artemisinin treatment conditions. Data shows the mean ±SEM for three independent experiments. (B) MA plot showing the deregulation in the expression of protein coding genes during artemisinin treatment with 30nM concentration for 6 hours and (C) during temperature stress at 40°C for 6 hours. (D) Expression profiles of the genes bound to PfGCN5 during stress conditions. PfGCN5 bound genes are upregulated upon stress induction as compared to control condition.

Further to investigate the role of PfGCN5 during stress conditions, we overexpressed HAT and bromo domains of PfGCN5 under normal and stress conditions as previous attempts failed to knockout GCN5 in *P. berghei* and *P. falciparum* indicating it is essential for parasite survival [43, 44]. Remarkably, we observed cell death upon overexpression of PfGCN5 HAT and bromo domains during stress conditions (S3C Fig). This in turn suggests that overexpression of PfGCN5 possibly leads to hyperactivation of stress responses, eventually resulting in cell death.

### PfGCN5 helps in maintenance of homeostasis during artemisinin treatment

Responses to oxidative stress and protein damage are shown to mediate emergence of artemisinin resistance in malaria parasites [15, 28, 29, 45, 46]. Interestingly, 775 new PfGCN5 bound sites were acquired under artemisinin stress conditions as indicated by ChIP-sequencing of PfGCN5 (Fig 4A). Moreover, gene ontology of newly acquired PfGCN5 bound genes under artemisinin stress conditions includes pathways such as ubiquitin-dependent protein catabolic process, cellular response to stimuli and response to drug, which are known to be deregulated in artemisinin resistant parasites (Fig 4B). Of interest is the binding of PfGCN5 at BiP and T-complex protein 1 (TCP1) ring (TRiC) chaperone genes. It is plausible that the higher expression of PfGCN5 upregulates BiP and TRiC chaperones, thus assisting the unfolded protein response in artemisinin resistant parasites [28]. This indicates that PfGCN5 might be playing an important role in the emergence of artemisinin resistance by regulating stress responsive pathways in *P. falciparum*.

**Figure 4:**
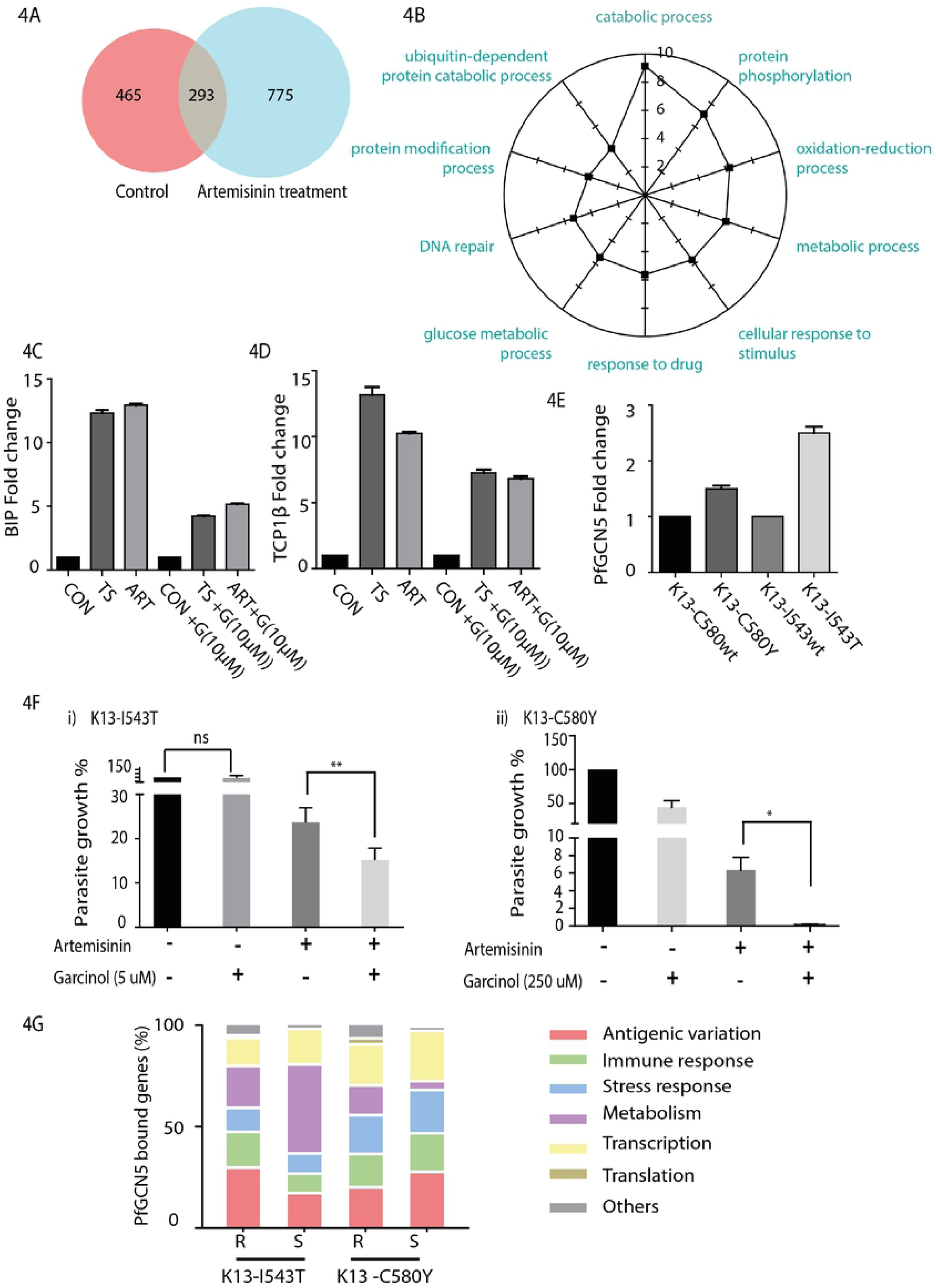
PfGCN5 shows prolific genomic binding during artemisinin treatment and in artemisinin resistant parasites. (A) Venn diagram showing the number of genes occupied by *Pf*GCN5 during normal conditions and during artemisinin treatment (30nM). (B) Gene ontology of the genes which are exclusively bound to *Pf*GCN5 during artemisinin treatment. PfGCN5 is found to be enriched on the genes which are known to be deregulated in artemisinin resistant parasites. This indicates the role of PfGCN5 during resistance generation. (C,D) RT-PCR results showing the upregulation of BiP and TCP1β during stress conditions and downregulation in the level of upregulation under garcinol treatment (10 µM) respectively. Data shows the mean ±SEM for three independent experiments. (E) Transcript level of expression of PfGCN5 in artemisinin resistant strains, K13-I543T (MRA-1241) and K13-C580Y (MRA-1236) in comparison to their sensitive counterparts, K13-I543wt (MRA-1253) and K13-C580wt (MRA-1254), respectively. (F) Change in the percentage parasite survival estimated through Ring Survival Assay (RSA) in presence of PfGCN5 inhibitor garcinol. i) K13-I543T (MRA-1241) parasites were treated with 5 µM garcinol and ii) K13-C580Y (MRA-1236) parasites were treated with 250 µM garcinol. Presence of garcinol decreases the artemisinin resistance in K13-I543T (MRA-1241) at a concentration which has otherwise no effect on parasite growth. Higher concentration of garcinol was used with a significant decrease in resistance level in K13-C580Y (MRA-1236) parasites. (G) Gene ontology enrichment of the genes which are bound by PfGCN5 in K13-I543T (MRA-1241), K13-C580Y (MRA-1236), K13-I543wt (MRA-1253), K13-C580wt (MRA-1254).

Furthermore, to understand the role of PfGCN5 in artemisinin drug resistance, we decided to use PfGCN5 inhibitor, garcinol. It is a specific inhibitor of PfGCN5 and showed an IC_50_ of ∼15 μM [47] (S4A-S4B Fig). We performed the quantitative RT-PCR during stress conditions both in presence and absence of garcinol (10µM). We found that both BiP and TCP1 β (T complex protein 1 subunit beta) were upregulated during the stress conditions (Fig 4C-4D). Interestingly, under garcinol treatment there is a decrease expression levels of BiP and TCP1 β under stress conditions (Fig 4C-4D).

Further to dissect the role of PfGCN5 in artemisinin drug resistance emergence and maintenance, we looked at transcript level of PfGCN5 in artemisinin resistant lines. Interestingly, PfGCN5 is upregulated in artemisinin resistant lines; K13-I543T (MRA-1241, RSA∼25%) and K13-C580Y (MRA-1236, RSA∼6%) by 2.5 and 1.5 fold than their sensitive counterparts, respectively (Fig 4E). We wondered if the inhibition of PfGCN5 activity resulted in change in drug sensitivity of the artemisinin resistant lines: K13-I543T and K13-C580Y. Ring survival assay (RSA) was performed in absence and presence of garcinol (used a concentration which has no or minimal effect on normal parasite growth). We observed 36.4% decreases in the level of resistance for K13-I543T artemisinin resistant line in the presence of PfGCN5 inhibitor, garcinol (Fig 4Fi). Interestingly, garcinol treatment of the artemisinin resistant parasites, K13-C580Y completely reverses the artemisinin resistance (Fig 4Fii) indicating that PfGCN5 plays an important role in artemisinin resistance.

In order to further understand the role of PfGCN5 in artemisinin resistance, we performed PfGCN5 ChIP sequencing in artemisinin resistant K13-I543T (MRA-1241) and K13-C580Y (MRA-1236) and their artemisinin sensitive counterpart K13-I543wt (MRA-1253) and K13-C580wt (MRA1254). We investigated the strain specific genes enriched for PfGCN5 binding and called their associated biological processes through the gene ontology analysis. Several biological processes were found to be conserved between the sensitive and resistant strain. These primarily include cellular adhesion, response to stimulus and antigenic variation, highlighting their regulation by PfGCN5 across strains (Fig 4G). Interestingly, a set of genes are uniquely enriched for PfGCN5 occupancy in the resistant strains. While, PfGCN5 is enriched on cellular metabolism and protein translation associated genes in K13-I543T strain, in K13-C580Y it is enriched on genes involved in vesicle fusion, and morphogenesis (Fig 4G). Deregulation of these biological pathways has been shown to be crucial for resistance acquisition in the field isolates of *P. falciparum* [30, 48]. Thus, our findings also reiterate an important aspect of resistance emergence posited earlier, that it is highly dynamic and can be shaped by independent underlying genetic and external environmental factors [14]. Together, these results suggest that PfGCN5 plays an important role in the regulation of stress responses, which are associated with drug resistance emergence.

### PfGCN5 interacts with PfAlba3 and regulates its chromatin binding

PfGCN5 has an unusually large N-terminal tail as compared to GCN5 in higher eukaryotes as well as other members of the phylum Apicomplexa. This large N-terminal region of PfGCN5 might play an additional role in protein-protein interactions to regulate *Plasmodium*-specific pathways. Further to understand the PfGCN5 mediated transcriptional regulation and to identify its interacting partners, we performed immunoprecipitation-coupled mass spectrometry using the two anti-PfGCN5 antibodies, α-HAT and α-peptide. We identified approximately 125 proteins interacting specifically with PfGCN5 (S3 Table), representing four major pathways namely chromatin assembly, response to stimuli, metabolic pathways and translation regulation (S5A Fig). Interestingly, one of the family of proteins identified as the interacting partners of PfGCN5 is PfAlba (Acetylation lowers binding affinity). PfAlbas are known to play diverse role during transcriptional and translational regulation [49, 50]. Alba proteins are also known to play important role in stress response pathways in higher eukaryotic system [51, 52]. As PfGCN5 was found to be majorly associated with stress responsive genes, we decided to further study PfGCN5 and PfAlba3 interaction.

First, to validate the interaction of PfGCN5 with PfAlba3, we cloned, expressed and purified recombinant His-tagged PfAlba3. As we were unable to express the full length PfGCN5 due to its large size, we cloned and overexpressed GST-tagged HAT and bromo domains of PfGCN5 (S1B Fig). Surprisingly, *in vitro* binding assay using recombinant His-tagged PfAlba3 and GST-tagged PfGCN5-HAT did not show any interaction (Fig 5A). Thus, it is possible that PfAlba3 either interacts with PfGCN5 outside of the HAT and bromodomain or it interacts indirectly with the PfGCN5 complex *in vivo*. Next, we performed immunoprecipitation using PfGCN5 peptide antibody and looked for PfAlba3 as its interacting partner in the pulled down fractions by Western blotting. As shown in Fig 5B, PfGCN5 co-elutes with PfAlba3 indicating an interaction with the PfGCN5 complex. Furthermore, immunofluorescence analysis suggested a partial colocalisation of PfGCN5 and PfAlba3 at trophozoite stages of *P. falciparum* (Fig 5C). Lastly, to understand the physiological role of PfGCN5 and PfAlba3 interaction, we performed *in vitro* acetyltransferase assays with the PfGCN5 complex and found that PfGCN5 indeed acetylates PfAlba3 (Fig 5D). Together, these data suggest that PfGCN5 interacts with PfAlba3 and mediates its acetylation.

**Figure 5:**
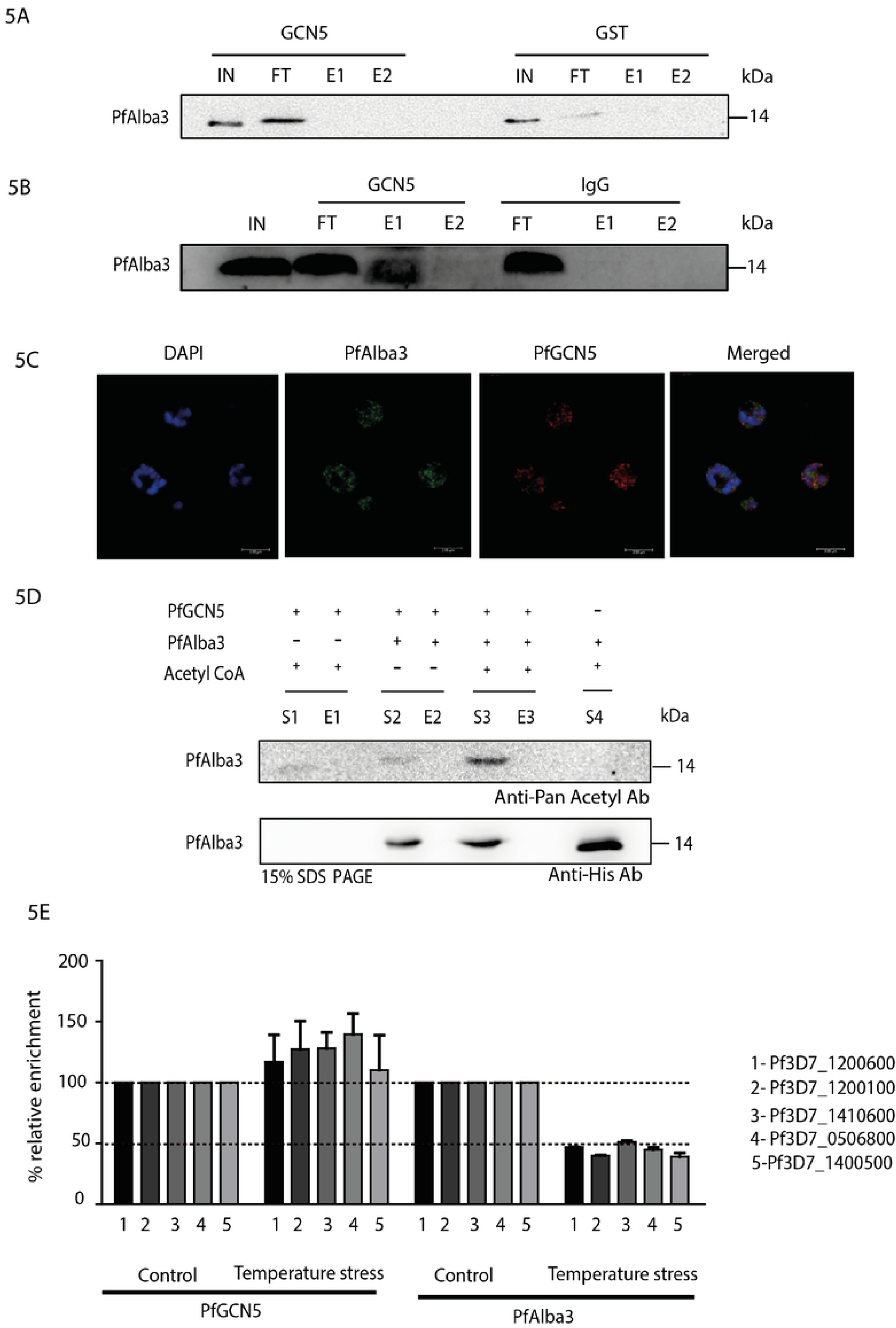
PfGCN5 regulates the binding of PfAlba3 to DNA through acetylation. (A) *In vitro* binding assay of recombinant PfGCN5 (HAT and bromodomain) and PfAlba3. Absence of binding was confirmed with Western blotting using Anti-His antibody. (B) Immunoprecipitation was performed to confirm the interaction of PfGCN5 and PfAlba3. PfGCN5 interacting proteins were pull down using PfGCN5 peptide antibody and binding was confirmed with Western blotting using PfAlba3 antibody. (C) Immunofluorescence assay was performed to check the localization of PfGCN5 and PfAlba3. Both the proteins were found to colocalize at certain regions indicating the positive interaction between them. PfGCN5 was visualized using Anti Rabbit Alexa 647 and PfAlba3 was labelled using Anti-Rat Alexa 488. Nucleus was stained using DAPI. Images were further processed for deconvolution using the Huygens Essential software. (D) Histone acetyltransferase (HAT) assay to verify PfGCN5 mediated acetylation of PfAlba3. The assay was performed using recombinant PfAlba3 and the PfGCN5 complex (pulled down with the help of PfGCN5 antibody). PfGCN5 was found to acetylate PfAlba3. (E) ChIP-qPCR showing the switching in the PfGCN5 and PfAlba3 enrichment on multivariant/stress responsive genes under temperature stress. ChIP was performed using PfGCN5 (peptide) and PfAlba3 antibodies.

Further to comprehend PfGCN5-mediated regulation of PfAlba3, we performed chromatin-immunoprecipitation using anti-PfGCN5 or anti-PfAlba3 antibodies. Previous studies [13] as well as our RNA sequencing data suggest that temperature stress results in deregulation of stress responsive and multicopy variant (var) genes. Importantly, virulence genes are known to be enriched for PfAlba occupancy and our data suggests enrichment of PfGCN5 (Fig 1D), hinting at involvement of the two factors in var expression regulation [49]. Thus, to explore the possible crosstalk between PfGCN5 and PfAlba3 in regulation of virulence and stress responsive genes, cells were subjected to temperature stress at 16 hpi for a period of 6 hours (5B Fig). This was followed by ChIP and qPCR with anti-PfGCN5 (peptide antibody) and anti-PfAlba3 (protein antibody) antibodies. Interestingly, we observed an increased occupancy of PfGCN5 and a corresponding decreased occupancy of PfAlba3 at stress responsive and virulence genes under temperature stress condition (Fig 5E). Thus, the interplay between PfGCN5 and PfAlba3 may play an important role in the regulation of various stress responsive and virulence genes depending on external cues.

## Discussion

### PfGCN5 is a stress/stimuli specific regulator and not a general transcriptional coactivator

*Plasmodium* must have evolved efficient machineries to overcome changes in environmental conditions experienced in two different hosts. The ability of *Plasmodium* to develop resistance against artemisinins is attributed to the competent stress responsive pathways and the unfolded protein response machinery, which are activated upon artemisinin exposure [14, 15]. Here, we establish the role of the histone acetyltransferase PfGCN5 as a global regulator of stress responsive pathways in *P. falciparum*. Genome-wide analysis of PfGCN5 occupancy shows that it is associated with stress responsive and multivariant gene family (virulence genes). Interestingly, PfGCN5 occupancy at various genomic loci was found to establish a transcriptionally poised state, which may allow these genes to be switched on or off immediately in response to stimuli. Such regulation is crucial for the genes implicated in stress response and host immune evasion. We and others have previously shown that H3K14ac, another histone modification mediated by GCN5, is specifically present on poised stress responsive genes in higher eukaryotic systems [53, 54] indicating a conserved role of GCN5 in *P. falciparum*. Together, these results suggest that PfGCN5 is not a general transcription coactivator and it specifically regulates the stress responsive and multicopy variant (virulence) genes in *P. falciparum*.

### PfGCN5 is an important modulator of artemisinin resistance

In order to get insights into the role of PfGCN5, we looked at the level of PfGCN5 transcript as well as genome wide binding sites during stress conditions i.e. heat stress and artemisinin exposure. We found that PfGCN5 is upregulated during stress conditions and its transcript level is comparable to artemisinin resistant parasite. Surprisingly, upon artemisinin treatment PfGCN5 is enriched on the genes important for the development of resistance against artemisinin. Corroborating PfGCN5 genome-wide binding with transcriptome data during stress conditions clearly indicates that PfGCN5 is associated with the genes which are upregulated during stress conditions (e.g. artemisinin exposure). Furthermore, upon interfering with the activity of PfGCN5 using its specific inhibitor, garcinol, we found a significant decrease in the level of artemisinin resistance in the K13-I543T mutant (MR4-1241, RSA-25%) and K13-C580Y (MRA-1236, RSA-6%). This in turn suggests that PfGCN5 is a global regulator of stress responsive genes, and plays an important role in artemisinin resistance maintenance.

Bhattacharjee *et al.* recently reported the amplified presence of PI3P vesicles which helps in mitigating the protein damage due to artemisinin treatment [48]. These vesicles house proteins like Kelch13, PfEMP1, BiP and others proteins required for maintaining homeostasis in artemisinin resistant parasite. Proteome analysis of these vesicles has revealed a list of proteins interacting with each other and possibly helping in emergence of artemisinin resistance [55]. PfGCN5 is one of the proteins detected in the vesicular proteome. We found a significant overlap in the proteins identified in the proteome analysis and PfGCN5 interacting partners (S6B Fig). In consonance, we also found various stress regulators such as heat shock proteins and Albas as interacting partners of PfGCN5. These interactions may play an important role in activation of stress response pathways upon artemisinin exposure. Moreover, PfGCN5 also regulates transcription regulation of BiP and T complex protein 1 beta subunit beta under stress conditions. Reports from higher eukaryotic systems have suggested that acetylation of BiP results in its dissociation from the protein kinase RNA-like endoplasmic reticulum kinase (PERK), which further results in phosphorylation of eIF2alpha leading to translation repression [56]. Moreover, we also found PfGCN5 to be enriched at the promoter of the Kelch13 gene, which possibly hints at its transcriptional regulation. Together, it suggests that PfGCN5 may play an important role in drug resistance generation either by directly regulating the expression of the genes important for emergence/maintenance of artemisinin resistance and/or by interacting with various key stress-regulators involved in resistance generation in *P. falciparum* (Fig 6).

**Figure 6:**
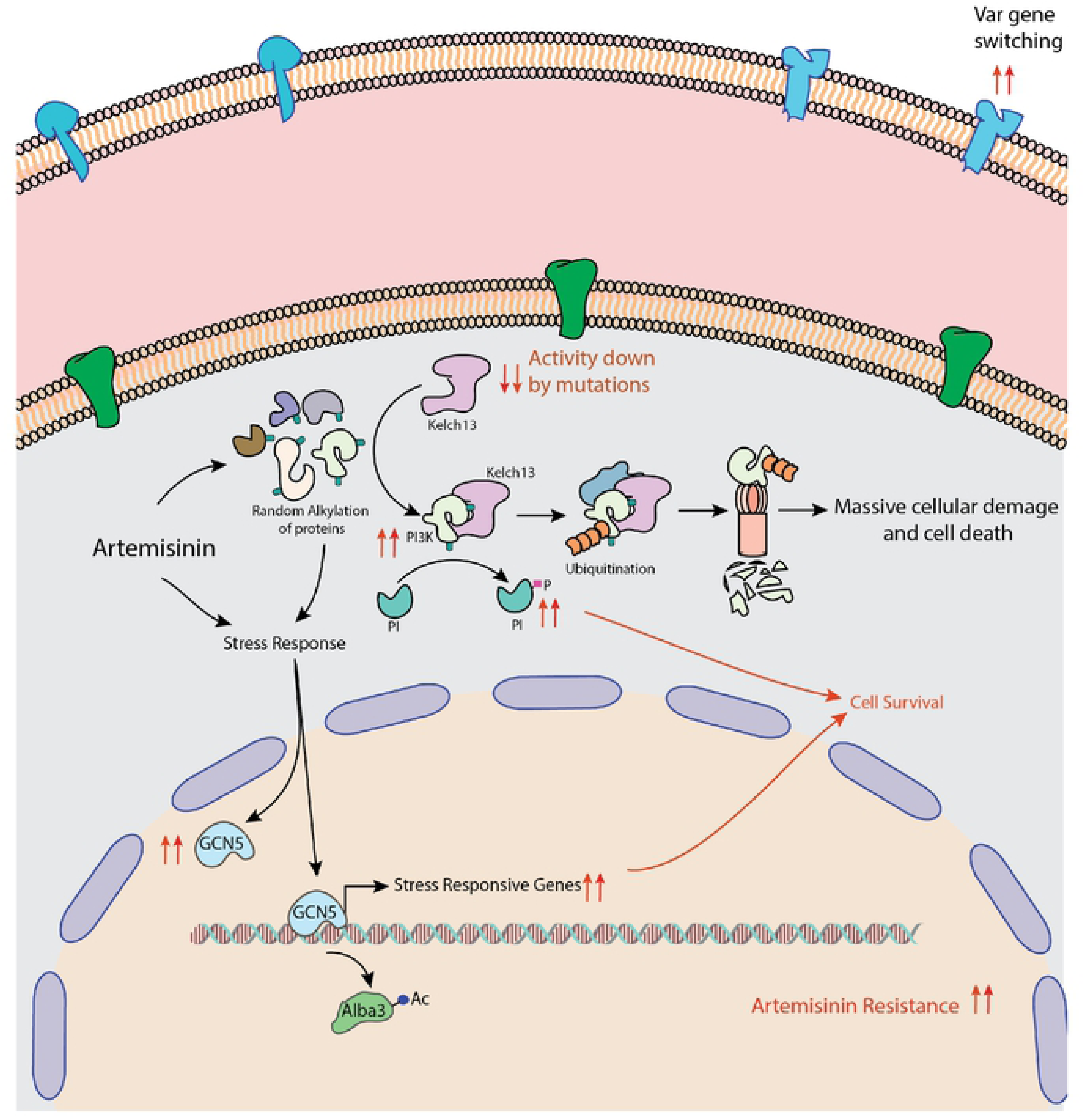
Mechanisms proposed for artemisinin resistance in *P. falciparum*. Model showing the role and interplay of Kelch13, PI3K and PfGCN5 in artemisinin resistance generation. Artemisinin treatment leads to random alkylation of proteins, which in turn are ubiquitinated by the protein ubiquitination complex (of which Kelch13 is an important ligase adapter component) and subjected to degradation by proteosomal degradation. One such protein is the PI3K (phosphatidylinositol 3 kinase), which is implicated in lipid metabolism and cell survival signaling. Massive alkylation by artemisinin exposure and/or oxidative stress activates PfGCN5 in the nucleus, which in turn upregulates the stress-responsive and unfolded protein response pathways. Thus, mutations in Kelch13 as well as upregulation of stress-responsive pathways by PfGCN5 help in artemisinin resistance generation.

### PfGCN5 regulates virulence gene expression upon stress induction

*Plasmodium falciparum* has evolved an extensive machinery to evade the host immune system through changes in the expression of multicopy variant proteins (var, rifin and stevor), which are expressed on the surface of infected RBCs [57, 58]. A switch in expression of these proteins also helps the parasite in evading splenic and immune clearance by a process called antigenic variation. Though the environmental cues responsible for virulence gene switching are not known, several factors have been identified to play regulatory roles in antigenic variation under physiological conditions [10, 57]. Various histone modifying enzymes like PfSir2, PfHda2, PfSET2 and PfSET10 are shown to repress expression of virulence genes [59–62]. *P. falciparum* heterochromatin protein 1 (HP1) is another key player known to repress the expression of virulence genes by binding to H3K9me2/3 and heterochromatinization [63, 64]. Here, we have identified PfGCN5 as a regulator of virulence gene expression switch under temperature stress condition.

Furthermore, we also found PfAlba3, a DNA/RNA binding protein, as an interacting partner of PfGCN5. PfAlba superfamily is known to play an important role in translation regulation in *P. falciparum* [50, 65]. Moreover, acetylation of PfAlba3 is known to lower its binding to DNA and results in gene activation [49]. Conversely, PfSir2a deacetylates PfAlba3 and makes it competent to bind DNA and leads to gene suppression [49]. Here we show that PfGCN5 binds to stress responsive and virulence genes and most probably regulates their expression by the acetylation of PfAlba3. Thus, the interplay between PfGCN5, PfSir2A and PfAlba3 possibly helps in regulation of stimuli dependent and virulence genes contributing to stress responsive and virulence phenotype.

Several studies in prokaryotes have investigated the link between virulence and resistance generation [66, 67]. There are clear evidences that virulence modulates resistance level in microorganisms and *vice versa* suggesting that there is a mechanism which tightly regulates both the processes. Geisinger *et al.* has showed the presence of a key stress response system in *Acinetobacter baumannii*, which enhances the virulence and resistance level in response to different physiological stresses [68]. Similarly, in *P. falciparum*, drug sensitivity of parasites is shown to be virulence dependent, where virulent parasites are shown to have higher likelihood to survive drug treatment [69]. Thus, it is plausible that regulation of both virulence as well stress responsive genes, which are responsible for drug resistance generation is mediated by same machinery involving GCN5 in *P. falciparum*.

Emergence of drug resistance against artemisinin is one of the biggest hurdles in malaria control and eradication. Recent reports have implicated stress responsive pathways in drug resistance generation (Fig 6). Understanding the regulation of stress responses and virulence gene expression is crucial to fathom the pathogenesis of the parasites. Our study identifies PfGCN5 as a global regulator of transcription of stress responses and virulence genes in *P. falciparum*. The outcome of this study could potentially be used to develop and screen inhibitors against drug resistant malaria parasites, which is one of the most prevalent parasitic diseases in the world.

## Experimental Procedures

### Parasite culture and transfection

*P. falciparum* strain 3D7 was cultured as previously described [70]. Briefly, parasites were cultured in RPMI1640 medium supplemented with 25 mM HEPES, 0.5 % AlbuMAX I, 1.77 mM sodium bicarbonate, 100 μM hypoxanthine and 12.5 μg ml^−1^ gentamicin sulfate at 37 °C. Parasites were sub-cultured after every two days. Subculturing was done by splitting the flask into multiple flasks in order to maintain parasitemia around 5%. Hematocrit was maintained to 1 - 1.5% by adding freshly washed O +ve human RBC isolated from healthy human donor. Synchronization was done with the help of 5% sorbitol in ring stage. Late stage synchronization was performed using the Percoll density gradient method (63%). Parasitemia was monitored using Giemsa staining of thin blood smear.

### Antibodies

Anti-actin (Sigma A2066) and Anti-Rabbit IgG (OSB PM035) were used for Western blotting and immunoprecipitation, respectively. Goat Anti-Rabbit Alexa Fluor 647 (A21245), Goat anti-Rat Alexa Fluor 488 (A 11006), Goat Anti-Rabbit Alexa Fluor 488 (A11034) were used for immunofluorescence. Rabbit polyclonal antibodies against PfAlba3 resulting from immunizations of rabbits with the KLH-conjugate peptide IGKRMFTGNEEKNP were obtained from GenScript Corporation [65]. Rat polyclonal antibodies against full-length recombinant GST-tagged PfAlba3 were from GenScript Corporation. For generating PfGCN5 peptide antibody, online software LBtope was used for selecting the antigenic peptide. PfGCN5 peptide (CEYCNVLYDGNELLRKRK) used for raising antibody was obtained from Apeptide Co., Ltd., China. PfGCN5 peptide was conjugated to Keyhole limpet hemocyanin (KLH) carrier protein for the immunization purpose. Both Anti-GCN5 peptide and protein antibodies were raised at The National Facility for Gene Function in Health and Disease, IISER Pune. The New Zealand White rabbits (3-4 months old) were used for antibody generation. Antibodies were further purified using affinity chromatography on the sulfolink resin.

### Western blotting

Parasites were harvested using 0.15% saponin. Parasites pellets were washed using phosphate buffer saline (PBS). Parasites were lysed using ice cold parasite lysis buffer (TRIS pH 8.0, 150 mM sodium chloride (NaCl), 0.5% nonyl phenoxypolyethoxylethanol (NP-40), 0.5% sodium deoxycholate, 0.1 mM ethylenediaminetetraacetic acid, 1.5 mM magnesium chloride (MgCl_2_), 1X protease inhibitor cocktail (PIC), 1 mM phenylmethylsulfonyl fluoride (PMSF). Three freeze thaw cycles were performed using liquid nitrogen to achieve proper lysis of the parasites. To get rid of debris, parasites were spun at 17949 x g for 30 minutes. Supernatant was transferred to another tube. The lysate proteins were separated on 7.5% - 12% polyacrylamide gels and transferred to PVDF membrane. The membrane was blocked using 5% skimmed milk and probed using primary antibody overnight at 4°C. After overnight incubation membrane were washed using 1X Tris-buffered saline, 0.1% Tween 20 (TBST) followed by 1hr incubation with secondary antibody in TBST (1:5000, Biorad). Three washes were given for 10 minutes each after the secondary antibody incubation. Blots were developed using Clarity Western ECL substrate (Biorad).

### Immunofluorescence Assay

Parasites were fixed using 4% PFA and 0.00075% glutaraldehyde for 30 minutes at 37°C. Permeabilisation was carried out using 0.1% Triton X-100 in PBS. Washing was performed using 1X PBS after every step. Blocking was done using 3% BSA for 1hr at room temperature followed by incubation with primary antibody in BSA for 3 hours. Three PBS washes were given to remove the unbound primary antibody. Secondary antibody incubation was done for 1 hour at room temperature. Parasites were washed before mounting on glass slides using ProLong Gold Antifade with DAPI (Invitrogen).

### Quantitative RT-PCR

RNA isolation was carried out using TRIzol reagent (Biorad). 2 μg of DNAse free RNA was used for cDNA synthesis using ImProm-II Reverse transcription system (Promega), as per the manufacture’s recommendation. Random primers were used for the cDNA synthesis. Real time PCR was carried out using CFX96 Real Time PCR detection system (Biorad). 18S rRNA and tRNA synthetase were used as an internal control to normalize for variability across different samples. Quantification of the expression was done with the help of fluorescence readout of SYBR green dye incorporation into the amplifying targets (Biorad). Each experiment included technical triplicates and was performed over three independent biological replicates. Primers details for the RT qPCR are given in S4 Table.

### Chromatin Immunoprecipitation

Infected RBCs were crosslinked using 1% formaldehyde (Thermo Scientific, 28908) for 10 mins at RT. 150 mM glycine was added for quenching the cross-linking reaction. The samples were washed using 1X PBS (chilled) before proceeding with lysis. Sample homogenization was performed using swelling buffer (25 mM Tris pH 7.9, 1.5 mM MgCl2, 10 mM KCL, 0.1% NP40, 1 mM DTT, 0.5 mM PMSF, 1x PIC) followed by cell lysis in sonication buffer (10 mM Tris–HCl pH 7.5, 200 mM NaCl, 1 % SDS, 4 % NP-40,1mM PMSF, 1X PIC). Sonication was performed using Covaris S220 to obtain the chromatin size of 200-400 bp. Pre-clearing was performed for 1 hour at 4°C using recombinant protein G conjugated sepharose beads with continuous gentle inverting. 30 μg purified chromatin was used per antibody (both α-HAT and α-peptide antibodies) and incubated for 12 h at 4°C. Samples were then incubated with saturated Protein G Sepharose beads for 4 hours at 4°C. Bound chromatin was finally washed and eluted using ChIP elution buffer (1 % SDS, 0.1 M sodium bicarbonate). Both IP sample and input were reverse crosslinked using 0.3M NaCl overnight at 65°C along with RNAse. Proteinase K treatment was performed at 42°C for 1 hour. Finally DNA was purified using phenol chloroform precipitation. Target sites identified from ChIP sequencing analysis were further validated by ChIP-qPCR using the Biorad SYBR Green Master Mix (Biorad). Primers details for the ChIP-qPCR are provided in S5 Table. Gene ontology was performed using PlasmoDB (www. plasmodb.org). Gene ontology terms along with number of genes in each category are given in S6 Table and S7 Table.

### ChIP-sequencing Library preparation and sequencing

ChIP-sequencing libraries for all the samples were prepared from 5-10 ng of DNA using the NEB Next Ultra II DNA Library Prep kit. Chromatin immunoprecipitated, fragmented DNA samples were end repaired and adapters ligated. Size selection was performed using Agencourt XP beads (Beckman Coulter). Adapter ligated fragments were PCR amplified using indexing primers followed by purification using the Agencourt XP beads (Beckman Coulter). The library electropherograms were assessed using Agilent Bioanalyzer 2100 and Agilent DNA 1000 kit. The libraries were pooled in equimolar concentration and 50 bp reads were sequenced using Illumina HiSeq2500 (BENCOS Research Solutions Pvt. Ltd., Maharashtra).

### Data pre-processing and peak calling

ChIP-seq data were mapped to *Plasmodium falciparum 3D7* genome version 37 (http://plasmodb.org/plasmo/) using Bowtie2 with default parameters. The mapped reads were used for peak calling against an input control data, using the MACS2 peak calling software (default parameters) [71]. Peaks were annotated using Bedtools [72]. ChIP-seq signals were background subtracted using MACS2 bdgcmp tool and the significantly enriched peaks were visualized using Integrative Genomics Viewer (IGV).

### Average profile calculations

We extracted the tag density in a 5 kb window surrounding the gene body using the seqMINER tool which generates heatmap as well as the enrichment profiles of factors over gene bodies [73]. For average gene profiles, genes (+/-5000 bp from binding site) were divided in 100 bins relative to the gene length. Moreover 10 equally sized (50 bp) bins were created on the 5’ and 3’ of the gene and ChIP-seq densities were collected for each dataset in each bin.

### Data source and analysis

ChIP seq data for Heterochromatin protein 1 (HP1) trophozoite stage was downloaded from Gene Expression Omnibus (http://www.ncbi.nlm.nih.gov/gds) with accession number: GSM2743113. Histone modification ChIP-seq data sets were downloaded from database under the accession number GSE63369. seqMINER [73] was used for generating scatter plots and average gene occupancy profiles. Correlation analysis and box plot were generated using ‘R’ software (http://r-project.org/).

### Stress induction

Parasites were subjected to heat and therapeutic (artemisinin treatment) stresses for 6 hours from late ring (∼17 hrs) to early trophozoite (∼23 hrs) stage. Double synchronization was carried out to achieve tight synchronization of parasite stages. Parasites were exposed to a) Heat stress (40°C for 6 hours) and b) Therapeutic stress (30 nM artemisinin for 6 hours).

### RNA sequencing and Data analysis

Parasites were harvested for RNA isolation after 6 hours of stress induction. Total RNA was isolated using TRIzol reagent according to the protocol. DNAse treated RNA was used for cDNA synthesis. Quality of the RNA was verified using Agilent Bioanalyzer 2100. Three biological replicates were pooled together for performing RNA sequencing. The cDNA libraries were prepared for samples using Illumina TruSeq RNA library preparation kit. Transcriptome sequencing was performed using Illumina NextSeq 500 system (1×150 bp read length) at BioServe Biotechnologies (India) Pvt Ltd. Hyderabad in replicate. Quality control of the RNA-sequencing reads was performed using FASTQC and reads were trimmed based on the quality estimates. The quality verified reads were then mapped onto the reference genome (PlasmoDB_v37) using the HISAT2 software (New Tuxedo Suite). After verification of the mapping percentage, the alignment data (SAM format) was converted into its binary counterpart (BAM format) using samtools. The same step also sorts the aligned reads positionally according to their genomic coordinates, making them easier to process further. In order to quantify the reads mapped onto the genomic features (genes, exons, etc.), the htseq-count feature was used. The count data was then used to perform differential gene expression (DGE) analysis and statistical validation using the Deseq2 package in the R computational environment. MA plot is generated using ‘R’ software (http://r-project.org/).

### Immunoprecipitation

In order to harvest the parasites, infected RBCs were lysed using 0.15% saponin at 37°C. Harvested parasites were then lysed using ice cold parasite lysis buffer (20 mM TRIS pH 8.0, 150 mM NaCl, 0.5% NP-40, 0.5% sodium deoxycholate, 0.1 mM EDTA, 1.5 mM MgCl_2_, 1X PIC, 1 mM PMSF). Lysed parasites were then centrifuged at 20817 x g for 30min at 4°C. Pre-clearing was performed using recombinant protein G conjugated sepharose beads for 1 hour at 4°C. Precleared lysate was then used for overnight incubation with antibody at 4°C. After the overnight incubation of lysate with antibody, sepharose Protein G beads were added to the lysate for 4 hours incubation. Washes were done using immunoprecipitation buffer (25 mM TRIS pH 7.9, 5 mM MgCl_2_, 10% glycerol, 100 mM KCl, 0.1% NP-40, 0.3 mM DTT) followed by elution of the proteins using glycine (pH – 2.5). Eluted proteins were neutralized using 1 M Tris pH 8.8. For mass spectrometry analysis samples were digested with trypsin for 16 hrs at 37°C. The digested samples were cleared using C18 silica cartridge. Peptides were then analysed using EASY-nLC 1000 system (Thermo Fisher Scientific) coupled to QExactive mass spectrometer (Thermo Fisher Scientific) equipped with nanoelectrospray ion source (Valerian Chem Private Limited, New Delhi). Immunoprecipitation followed by mass spectrometry was performed in three biological replicates. Samples were processed and RAW files generated were analyzed with Proteome Discoverer against the Uniprot P.falci reference proteome database. For Sequest search, the precursor and fragment mass tolerances were set at 10 ppm and 0.5 Da, respectively. The protease used to generate peptides, i.e. enzyme specificity was set for trypsin/P (cleavage at the C terminus of “K/R: unless followed by “P”) along with maximum missed cleavages value of two. Carbamidomethyl on cysteine as fixed modification and oxidation of methionine and N-terminal acetylation were considered as variable modifications for database search. Both peptide spectrum match and protein false discovery rate were set to 0.01 FDR.

### Protein expression and purification

PfGCN5 (HAT and bromodomain) DNA sequence was amplified from parasite genomic DNA using gene specific primers. The PCR-amplified fragment was cloned in frame with glutathione S-transferase (GST) fusion protein in pGEX-4T1 plasmid vector using XhoI and BamHI restriction enzymes. For expression in *E.coli*, pGEX-4T1 (GCN5) plasmid was transformed in BL21 (DE3) star competent cells. Expression was induced at an optical density of 0.6 at 600 nm, with 0.5 mM isopropyl-1-thio-β-d-galactopyranoside (IPTG) for 5 hrs at 25°C. Protein was purified using glutathione sepharose 4B beads (GE healthcare life science). 20 mM concentration of reduced glutathione was used for protein elution. PfAlba3 was cloned in pET28a^+^ vector using NdeI and XhoI. Histidine-tagged PfAlba3 was expressed in the *E. coli* BL21 (DE3) competent cells. Expression was induced at an optical density of 0.6 at 600nm, with 0.5 mM IPTG for 5 hours at 25°C. Protein purification was performed using Ni-NTA beads. Protein was eluted using different concentration of Imidazole. Purified proteins were dialyzed and stored at −20°C. Primers details for the cloning are provided in S8 Table.

### *In vitro* interaction assay

*In vitro* interaction study was carried out using GST-tagged PfGCN5 and His tagged Alba3 proteins. Recombinant proteins (2ug) were incubated together overnight at 4°C. GST protein was used as the negative control. Glutathione beads were added to the protein mix for 4 hours at 4°C. The beads were washed and the bound proteins were eluted from the beads using 20 mM reduced glutathione. Western blotting was performed using Anti-His antibody to verify presence of PfAlba3 in the elutions.

### Ring stage survival assay (RSA)

*In vitro* RSA was performed according to the protocol described in Witkowski *et al.* (2013) [74]. Parasites were synchronized at early ring stage. Tightly synchronised 0-3 hrs rings were given 700 nM of artemisinin for 6 hrs. Drug was washed after 6 hrs with RPMI. Culture was then cultivated for 66 hrs. Parasites were then lysed and the parasite growth was calculated with the help of SYBR green I reagent which intercalates with the DNA and gives a fluorescent readout upon excitation. Parasite survival rate was calculated comparing the growth between drug treated and untreated control.

### Data access

ChIP-sequencing data for PfGCN5 as well as gene expression data (RNA sequencing) for different conditions are submitted to Sequence Read Archive (SRA) under ID SUB5640877.

### Ethics Statement

This study does not involve human participants. Human RBCs used in this study were obtained from the KEM Blood Bank (Pune, India) as blood from anonymized donors. Approval to use this material for *P. falciparum in vitro* culture has been granted by the Institutional Biosafety Committee of Indian Institute of Science Education and Research Pune (BT/BS/17/582/2014-PID). The use of rabbits in this study for immunization (IISER/IAEC/2017-01/008) was reviewed and approved by Indian Institute of Science Education and Research (IISER)-Pune Animal House Facility (IISER: Reg No. 1496/GO/ReBi/S/11/CPCSEA). The approval is as per the guidelines issued by Committee for the Purpose of Control and Supervision of Experiments on Animals (CPCSEA), Govt. of India.

## Acknowledgements

The following reagent was obtained through BEI Resources (www.mr4.org), NIAID, NIH: *Plasmodium falciparum*, Strain IPC 3445 (MRA-1236), Strain IPC 4912 (MRA-1241), contributed by Didier Menard and *Plasmodium falciparum*, Strain Cam2_rev (MRA-1254), Strain IPC Cam2_rev (MRA-1253), contributed by David A. Fidock. We thank Dr. Artur Scherf from Institut Pasteur, Paris, France, for his support. This work was supported by grants under DST INSPIRE (IFA-13, LMBM-53) and DBT-IYBA (BT/08/IYBA/2014-17) from Government of India to KK. MR is supported by DBT-SRF fellowship. MR is also supported by EMBO short term fellowship to carry out part of this work. The funders had no role in study design, data collection and analysis, decision to publish, or preparation of the manuscript.

## Supporting Information Figure Legends

**Supplementary Table S1. PfGCN5 bound sites identified using ChIP sequencing.** ChIP sequencing of PfGCN5 using the α-peptide antibody was performed during early trophozoites. The sites identified to be bound by PfGCN5 are listed in the table according to their decreasing fold enrichment.

**Supplementary Table S2. Genes deregulated during stress conditions.** RNA sequencing was performed during stress conditions to identify the genes deregulated. List of genes along with their tag count is listed in table 2.

**Supplementary Table S3. PfGCN5 interacting proteins.** PfGCN5 interacting proteins were identified using the both PfGCN5 α-HAT and α-peptide antibody. List of proteins which were found using both these antibodies are mentioned in the table 3.

**Supplementary Table S4. Primers used in the study**. Sequences of the RT-PCR primers used in the study.

**Supplementary Table S5. Primers used in the study.** Sequences of the quantitative PCR primers used in this study.

**Supplementary Table S6. Genes identified in each gene ontology term.** Gene ontology of the genes bound with PfGCN5 was performed using Plasmodb. Number of genes which were found in each category of the gene ontology term is mentioned in the table.

**Supplementary Table S7. Genes identified in each gene ontology term**. Gene ontology of the genes bound with PfGCN5 exclusively during artemisinin treatment was performed using Plasmodb. Number of genes which were found in each category of the gene ontology term is mentioned in the table.

**Supplementary Table S8. Primers used for the cloning of PfGCN5.** Sequences of the primers used for cloning of PfGCN5 for protein expression and overexpression.

## Supplementary Figures

**Supplementary Figure S1:**
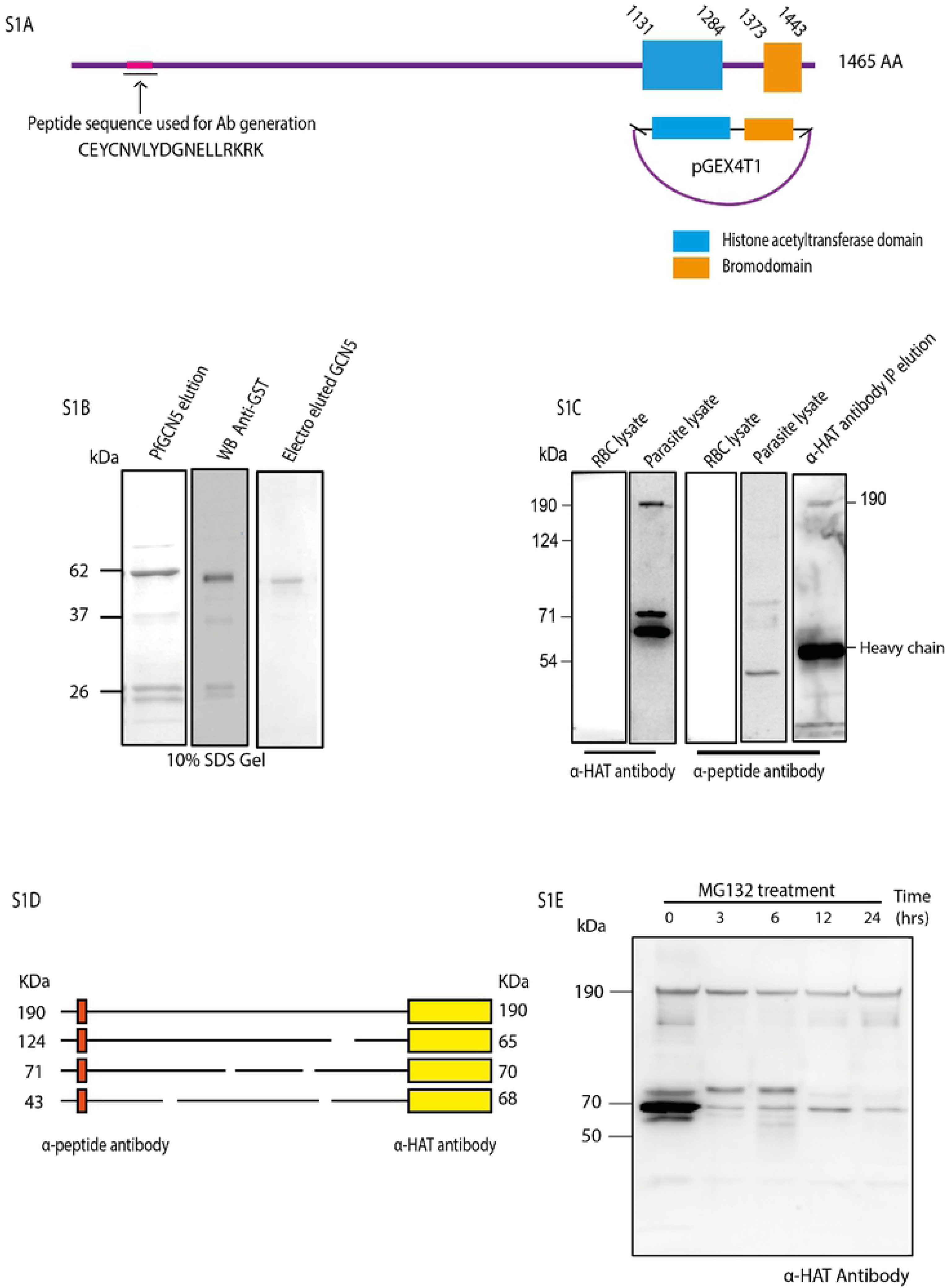
*Pf*GCN5 specific antibody generation. (S1A) Schematic diagram showing the domain organization of PfGCN5. Histone acetyltransferase (HAT) domain and bromodomain (represented in blue and orange colour, respectively) are present at C terminal end. PfGCN5 peptide from N-terminal region of the protein was commercially synthesized for raising antibody. The HAT and bromodomain was cloned in pGEX 4T1 vector for expression of PfGCN5 protein tagged with GST. Protein expression was induced using 0.5mM IPTG in BL21 (DE3) star competent cells. (S1B) Protein was purified using glutathione beads and eluted using reduced glutathione (20mM). Protein expression was confirmed using anti-GST Western blotting. Protein was further purified using electro elution before injected in rabbit for Anti-PfGCN5 antibody generation. Single band protein (PfGCN5-GST) was observed after electro elution in SDS-PAGE. Antiserum raised against PfGCN5 protein (HAT and bromodomain) was checked for specificity using bacterial lysate expressing PfGCN5. (S1C) Specificity of the antibody was further checked using parasite protein lysate from asynchronous culture. Western blotting result indicates the presence of more than one forms of PfGCN5 in *Plasmodium.* Western blotting was performed on proteins which are pulled down by α-HAT antibody and probed with α-peptide antibody. The presence of full length band indicates that both antibodies detect full length PfGCN5. (S1D) Schematic to possibly explain the bands which are observed during Western blotting for both the antibodies generated again PfGCN5. (S1E) In order to check whether the extra bands observed in Western blotting are the result of possible proteosomal degradation, parasites were treated with MG132 inhibitor for different duration of time. MG132 treatment resulted in significant decrease in number of bands observed. Only two bands were observed after 24hr treatment of MG132 which might indicate two isoforms of PfGCN5.

**Supplementary Figure S2:**
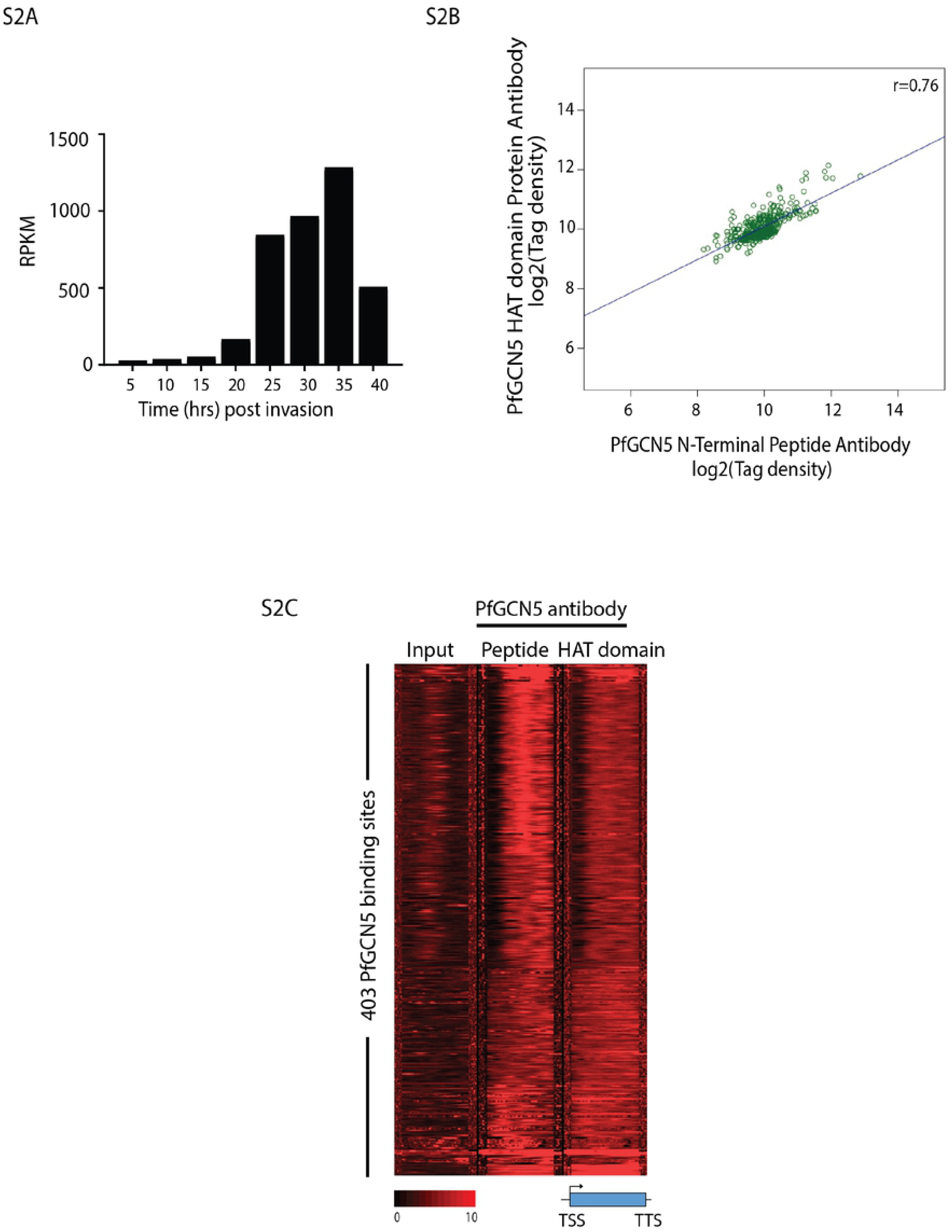
PfGCN5 is associated with virulence and stimuli induced genes. (S2A) Dynamics of PfGCN5 transcript expression during different stages of intraerythrocytic life cycle of *P.falciparum.* Expression profile suggests the low expression of PfGCN5 during the ring stages and a sudden burst of PfGCN5 mRNA expression during early trophozoite stage. (S2B) Scatter plot depicting a linear correlation of the target tag densities in ChIP pulldown performed using the scores of overlapping peaks from the PfGCN5 HAT domain and peptide antibodies. This is indicative of the fact that both antibodies have similar pulldown profile in ChIP sequencing reads. (S2C) Heat map showing PfGCN5 occupancy over 403 genes identified as the targets using PfGCN5 (HAT) antibody and peptide antibody.

**Supplementary Figure S3:**
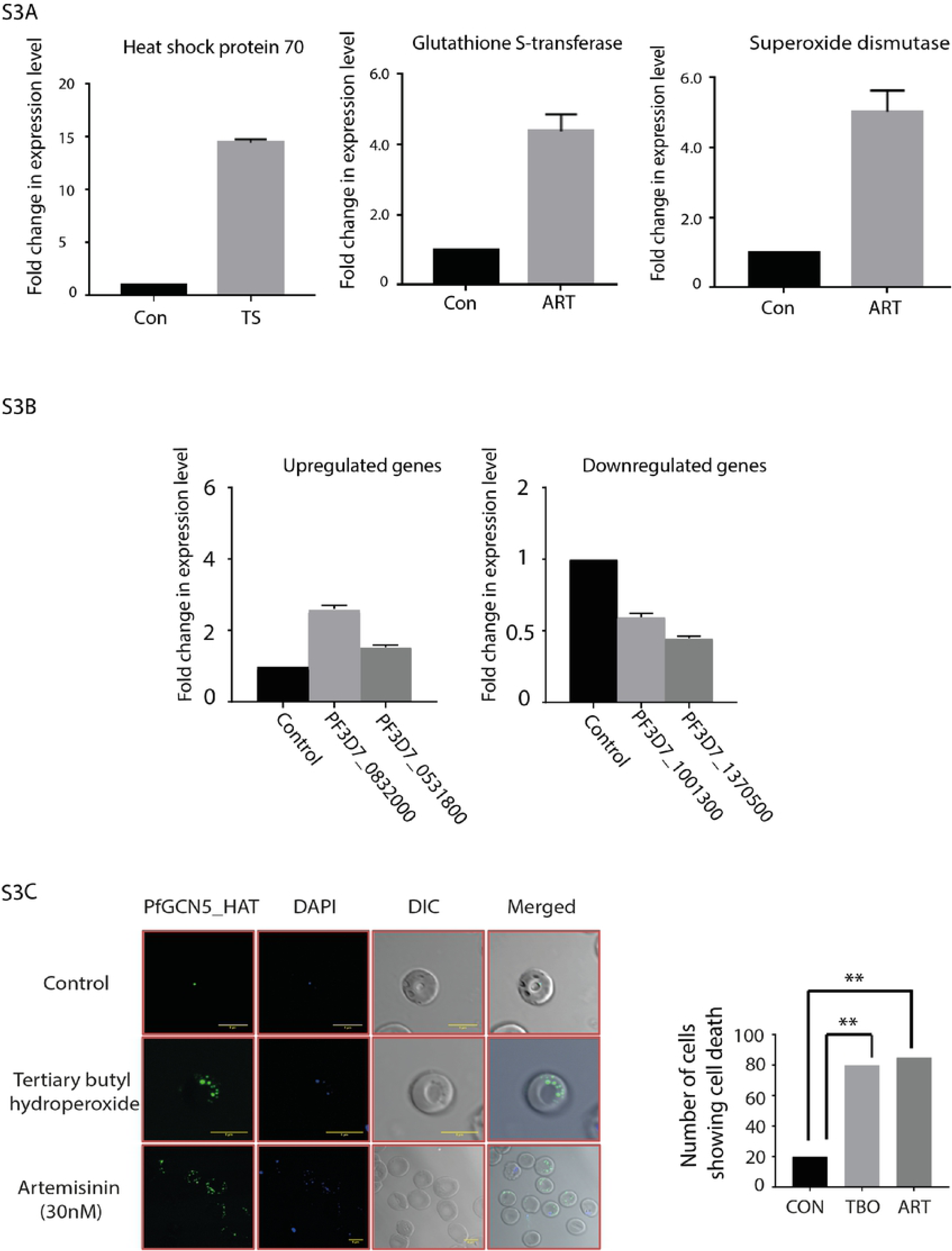
Differential gene expression during stress conditions. (S3A) Different markers genes were found to be deregulated during stress conditions. Temperature stress results in up regulation of HSP70. Similarly artemisinin (ART) treatment results in the increase in expression of Glutathione S-transferase and Superoxide dismutase which indicates the presence of ROS in parasites due to artemisinin treatment. Up regulation of these markers genes is indicative of the fact that stress is induced in the parasite upon artemisinin treatment and increase in temperature. (S3B) RT-qPCR validation of the genes which are deregulated during stress conditions, identified through RNA sequencing. (S3C) Parasites with episomal overexpression of truncated PfGCN5 (HAT and bromodomain) exhibited cell death during stress conditions. Artemisinin (Art, 30nM) and Tert-Butyl hydroperoxide (TBO, 10mM) were given to parasites for 6 hrs. TBO was used to induce ROS stress in parasites.

**Supplementary Figure S4:**
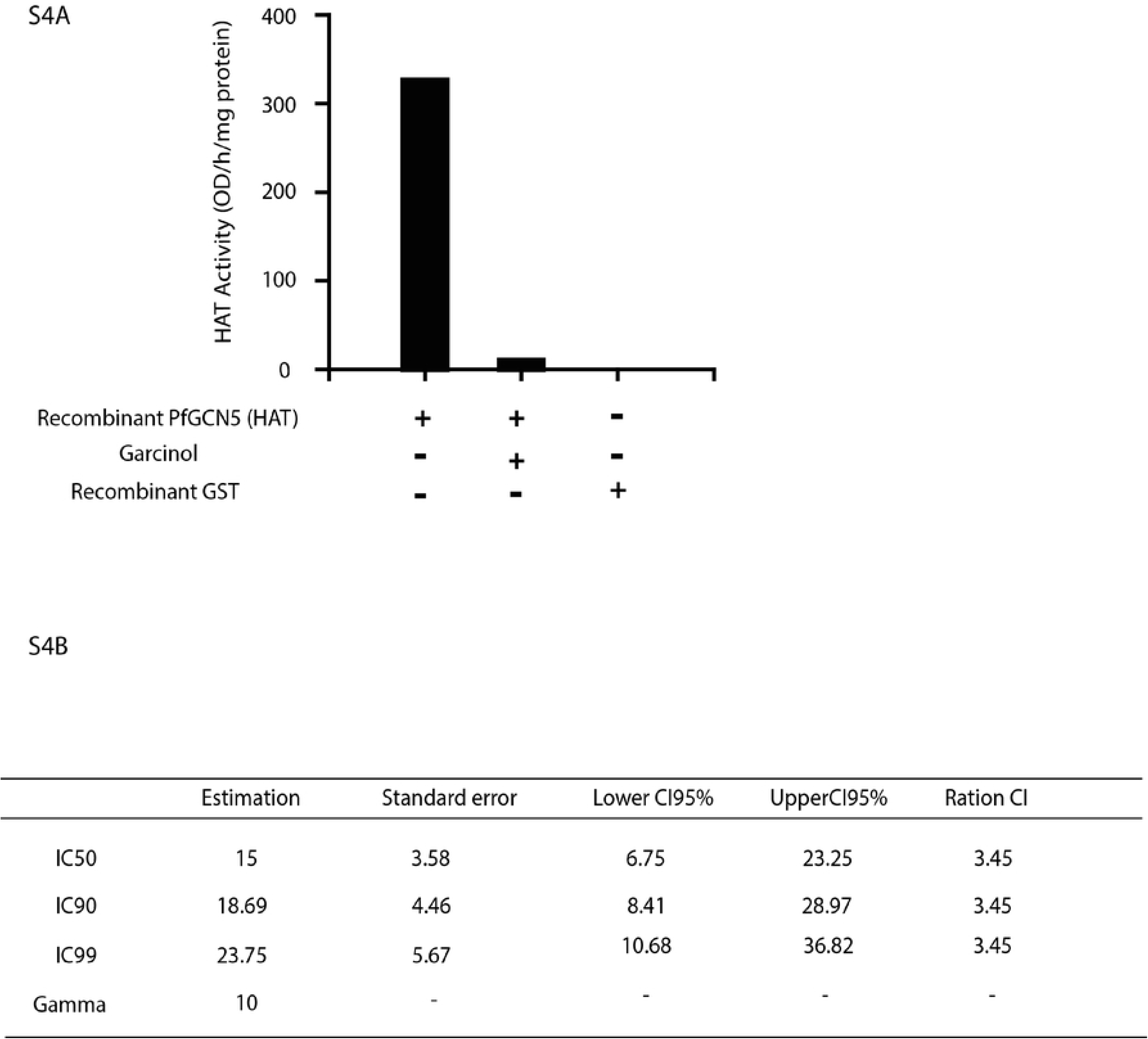
Recombinant HAT domain of PfGCN5 is catalytically active and can be inhibited by garcinol treatment. (S4A) Inhibition of histone acetylation activity of purified recombinant HAT domain of PfGCN5. 10 µM of Garcinol inhibits PfGCN5 HAT activity completely. (S4B) IC50 calculation of garcinol using dose response assay carried out over a period of 48 hours. The growth inhibition was measured using the SYBR green dye.

**Supplementary Figure S5:**
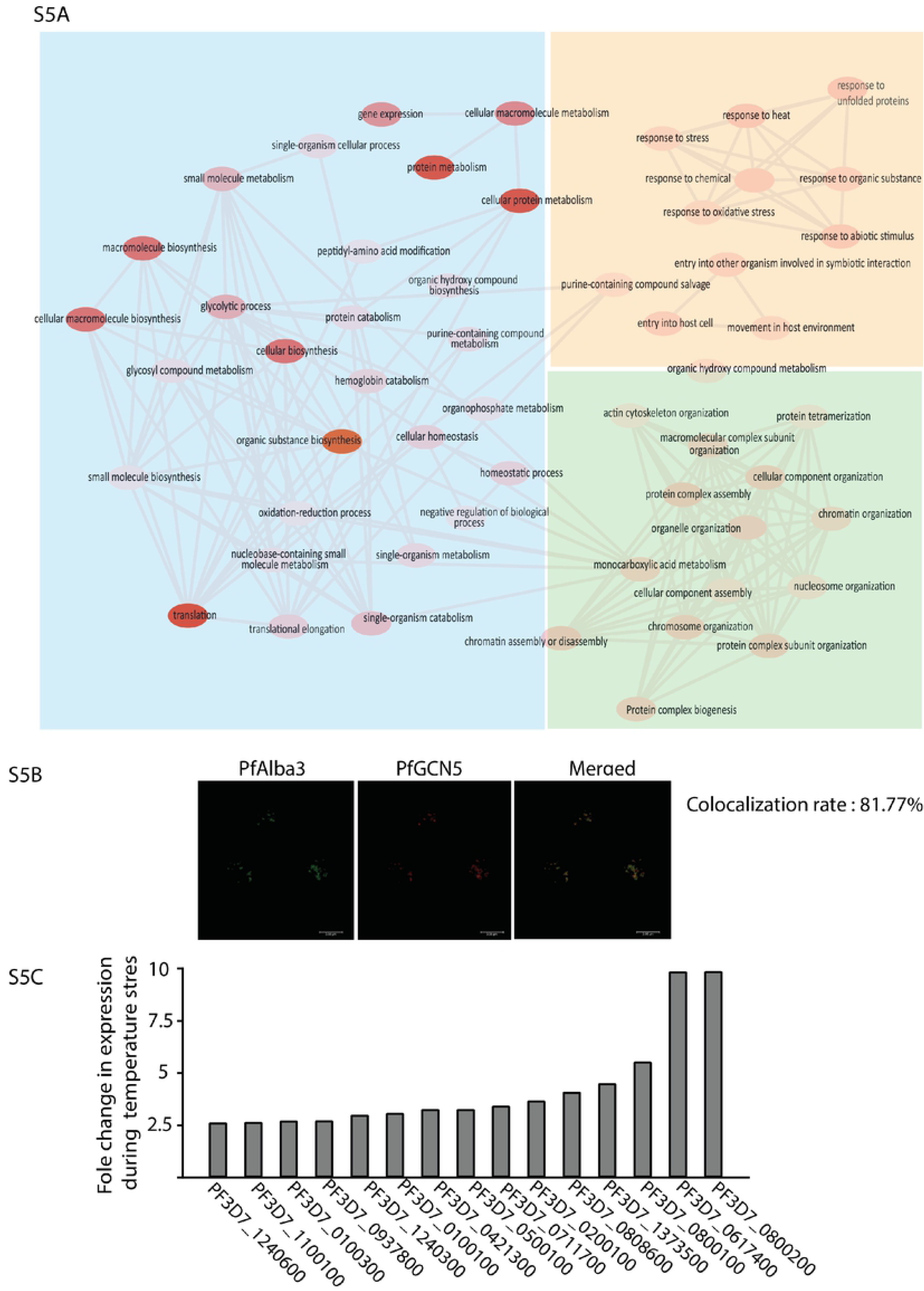
Gene Ontology of the protein interactors of PfGCN5; validation of interaction of PfGCN5-PfAlba3 and investigating the expression of PfAlba3 targets in stress conditions. (S5A) Gene ontology analysis of PfGCN5 interacting proteins indicate the overrepresentation of four major biological pathways namely chromatin assembly, response to stimuli, metabolic pathways and translation regulation. Gene ontology was performed using PlasmoDB and the plot was generated using Revigo (http://revigo.irb.hr/) (S5B) Immunofluorescence images to investigate the colocalisation of PfGCN5 and PfAlba3 show high rate for colocalisation, suggesting significant overlap between the two. (S5C) Change in the expression level of the various virulence genes during temperature stress. Temperature stress results in upregulation of more than one virulence genes in *Plasmodium*.

**Supplementary Figure S6:**
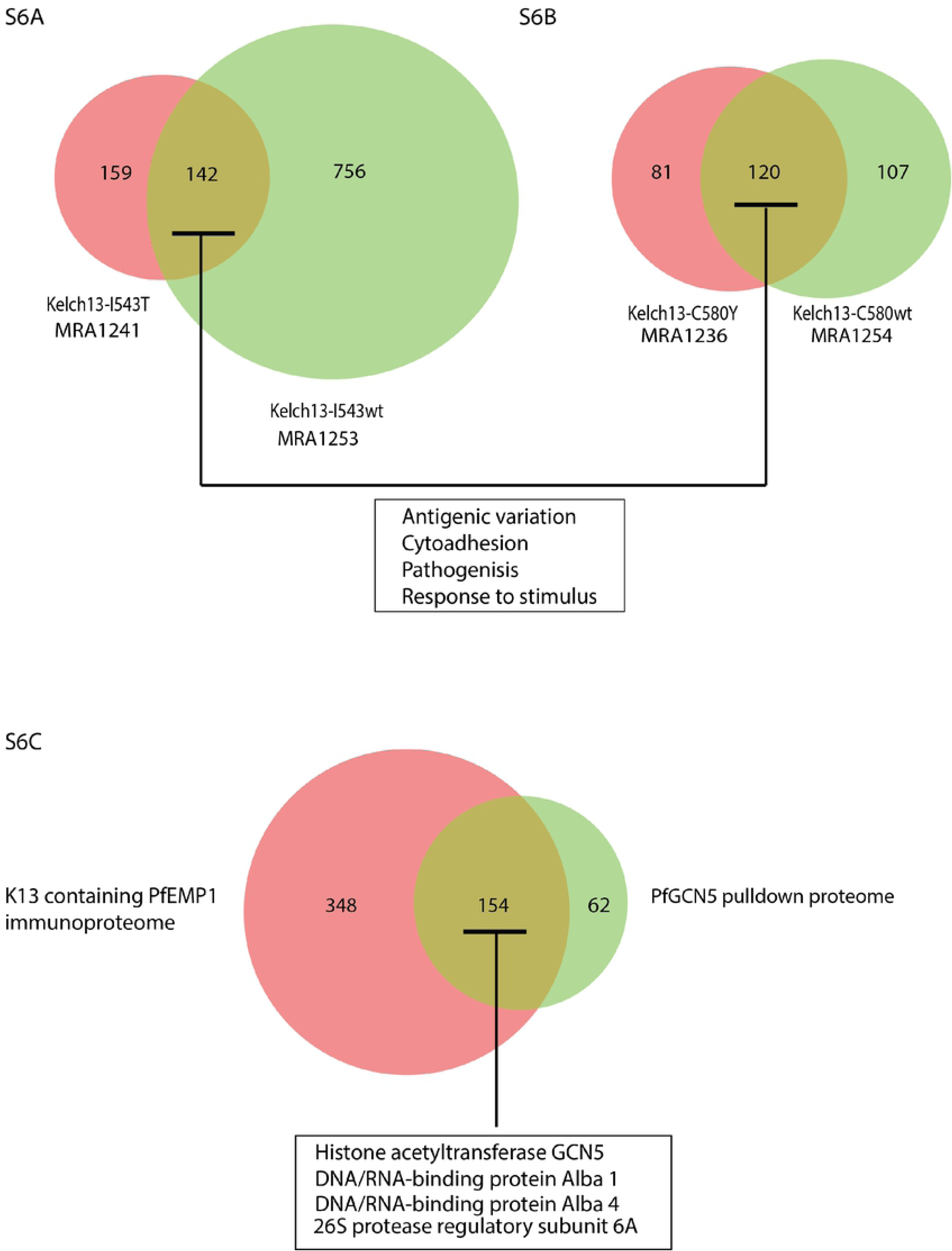
PfGCN5 regulates antigenic variation and stress response machinery in artemisinin resistant strains of *Plasmodium*. (S6A) Venn diagram showing the genes bound by PfGCN5 in K13-I543T (MRA-1241) and K13-C580Y (MRA-1236) mutant lines and their sensitive counterpart K13-I543wt (MRA-1253) and K13-C580wt (MRA-1254) respectively. Genes associated with antigenic variation and cellular adhesions were found to be core targets of PfGCN5 in both sensitive and resistant strains. Unique set of genes implicated in resistance in the field were enriched for PfGCN5 occupancy in the two resistant strains. (S6B) Venn-diagram showing the overlap between the various proteins which were pulled down during PfGCN5 immunoprecipitation and the protein found in the vesicles secreted by the artemisinin resistant parasites.

## References

1. WHO (2018) World Malaria Report.

2. Mamoun, C. B., Gluzman, I.Y., Hott, C., MacMillan, S.K., Amarakone, A.S., Anderson, D.L., Carlton, J.M., Dame, J.B., Chakrabarti, D., Martin, R.K., Brownstein, B.H., Goldberg, D.E. (2001) Co-ordinated programme of gene expression during asexual intraerythrocytic development of the human malaria parasite Plasmodium falciparum revealed by microarray analysis, Molecular microbiology. 39, 26–36.

3. Coleman, B. I. & Duraisingh, M. T. (2008) Transcriptional control and gene silencing in Plasmodium falciparum, Cell Microbiol. 10, 1935–46.

4. Cui, L., Lindner, S. & Miao, J. (2015) Translational regulation during stage transitions in malaria parasites, Annals of the New York Academy of Sciences. 1342, 1–9.

5. Karmodiya, K., Pradhan, S. J., Joshi, B., Jangid, R., Reddy, P. C. & Galande, S. (2015) A comprehensive epigenome map of Plasmodium falciparum reveals unique mechanisms of transcriptional regulation and identifies H3K36me2 as a global mark of gene suppression, Epigenetics & chromatin. 8, 32.

6. Duffy, M. F., Selvarajah, S. A., Josling, G. A. & Petter, M. (2013) Epigenetic regulation of the Plasmodium falciparum genome, Briefings in functional genomics. 13, 203–216.

7. Volz, J. C., Bartfai, R., Petter, M., Langer, C., Josling, G. A., Tsuboi, T., Schwach, F., Baum, J., Rayner, J. C., Stunnenberg, H. G., Duffy, M. F. & Cowman, A. F. (2012) PfSET10, a Plasmodium falciparum methyltransferase, maintains the active var gene in a poised state during parasite division, Cell Host Microbe. 11, 7–18.

8. Guizetti, J. & Scherf, A. (2013) Silence, activate, poise and switch! Mechanisms of antigenic variation in Plasmodium falciparum, Cell Microbiol. 15, 718–26.

9. Percário, S., Moreira, D. R., Gomes, B. A. Q., Ferreira, M. E. S., Gonçalves, A. C. M., Paula S. O. C. Laurindo, P. S. O. C., Vilhena, T. C., Dolabela, M. F. & Green, M. D. (2012) Oxidative Stress in Malaria, International Journal of Molecular Sciences. 13, 16346–16372.

10. Rosenberg, E., Ben-Shmuel, A., Shalev, O., Sinay, R., Cowman, A. & Pollack, Y. (2009) Differential, positional-dependent transcriptional response of antigenic variation (var) genes to biological stress in Plasmodium falciparum, PLoS One. 4, e6991.

11. Beckera, K., Tilley, L., Vennerstrom, J. L., Roberts, D., Rogerson, S. & Ginsburg, H. (2004) Oxidative stress in malaria parasite-infected erythrocytes: host–parasite interactions, International Journal for Parasitology 34, 163–189.

12. Engelbrecht, D. & Coetzer, T. L. (2013) Turning up the heat: heat stress induces markers of programmed cell death in Plasmodium falciparum in vitro, Cell Death Dis. 4, e971.

13. Oakley, M. S., Kumar, S., Anantharaman, V., Zheng, H., Mahajan, B., Haynes, J. D., Moch, J. K., Fairhurst, R., McCutchan, T. F. & Aravind, L. (2007) Molecular factors and biochemical pathways induced by febrile temperature in intraerythrocytic Plasmodium falciparum parasites, Infect Immun. 75, 2012–25.

14. Rocamora, F., Zhu, L., Liong, K. Y., Dondorp, A., Miotto, O., Mok, S., Bozdech, Z. Oxidative stress and protein damage responses mediate artemisinin resistance in malaria parasites, PLoS Pathog, e1006930.

15. Dogovski, C., Xie, S. C., Burgio, G., Bridgford, J., Mok, S., McCaw, J. M., Chotivanich, K., Kenny, S., Gnadig, N., Straimer, J., Bozdech, Z., Fidock, D. A., Simpson, J. A., Dondorp, A. M., Foote, S., Klonis, N. & Tilley, L. (2015) Targeting the cell stress response of Plasmodium falciparum to overcome artemisinin resistance, PLoS Biol. 13, e1002132.

16. Wellems, T. E., Panton, L. J., Gluzman, I. Y., do Rosario, V. E., Gwadz, R. W., Walker-Jonah, A. & Krogstad, D. J. (1990) Chloroquine resistance not linked to mdr-like genes in a Plasmodium falciparum cross, Nature. 345, 253–255.

17. Basco, L. K., Tahar, K. & Ringwald, P. (1998) Molecular Basis of In Vivo Resistance to Sulfadoxine-Pyrimethamine in African Adult Patients Infected with Plasmodium falciparum Malaria Parasites, ANTIMICROBIAL AGENTS AND CHEMOTHERAPY. 42, 1811–1814.

18. Nosten, F., van Vugt, M., Price, R., Luxemburger, C., Thway, K. L., Brockman, A., McGready, R., ter Kuile, F., Looareesuwan, S. & White, N. J. (2000) Effects of artesunate-mefloquine combination on incidence of Plasmodium falciparum malaria and mefloquine resistance in western Thailand: a prospective study, The Lancet. 356, 297–302.

19. Ridley, R. G. (2002) Medical need, scientific opportunity and the drive for antimalarial drugs, Nature. 415, 686–693.

20. Blasco, B., Leroy, D. & Fidock, D. A. (2017) Antimalarial drug resistance: linking Plasmodium falciparum parasite biology to the clinic, Nat Med. 23, 917–928.

21. Dondorp, A. M., Nosten, F., Yi, P., Das, D., Phyo, A. P., Tarning, J., Lwin, K. M., Ariey, F., Hanpithakpong, W., Lee, S. J., Ringwald, P., Silamut, K., Imwong, M., Chotivanich, K., Lim, P., Herdman, T., An, S. S., Yeung, S., Singhasivanon, P., Day, N. P., Lindegardh, N., Socheat, D. & White, N. J. (2009) Artemisinin resistance in Plasmodium falciparum malaria, N Engl J Med. 361, 455–67.

22. Fairhurst, R. M. & Dondorp, A. M. (2016) Artemisinin-Resistant Plasmodium falciparum Malaria, Microbiol Spectr. 4.

23. Hanboonkunupakarn, B. & White, N. J. (2016) The threat of artemisinin resistant malaria in Southeast Asia, Travel Med Infect Dis. 14, 548–550.

24. Mishra, N., Prajapati, S. K., Kaitholia, K., Bharti, R. S., Srivastava, B., Phookan, S., Anvikar, A. R., Dev, V., Sonal, G. S., Dhariwal, A. C., White, N. J. & Valecha, N. (2015) Surveillance of artemisinin resistance in Plasmodium falciparum in India using the kelch13 molecular marker, Antimicrobial agents and chemotherapy. 59, 2548–53.

25. Das, S., Manna, S., Saha, B., Hati, A. K. & Roy, S. (2018) Novel pfkelch13 gene polymorphism associates with artemisinin resistance in eastern India, Clinical infectious diseases : an official publication of the Infectious Diseases Society of America.

26. Mok, S., Imwong, M., Mackinnon, M. J., Sim, J., Ramadoss, R., Yi, P., Mayxay, M., Chotivanich, K., Liong, K.-Y. & Russell, B. (2011) Artemisinin resistance in Plasmodium falciparum is associated with an altered temporal pattern of transcription, BMC genomics. 12, 391.

27. Phompradit, P., Chaijaroenkul, W. & Na-Bangchang, K. (2017) Cellular mechanisms of action and resistance of Plasmodium falciparum to artemisinin, Parasitol Res. 116, 3331–3339.

28. Suresh, N., Haldar, K. (2018) Mechanisms of artemisinin resistance in Plasmodium falciparum malaria, Current Opinion in Pharmacology, 46–54.

29. Mbengue, A., Bhattacharjee, S., Pandharkar, T., Liu, H., Estiu, G., Stahelin, R. V., Rizk, S. S., Njimoh, D. L., Ryan, Y., Chotivanich, K., Nguon, C., Ghorbal, M., Lopez-Rubio, J. J., Pfrender, M., Emrich, S., Mohandas, N., Dondorp, A. M., Wiest, O. & Haldar, K. (2015) A molecular mechanism of artemisinin resistance in Plasmodium falciparum malaria, Nature. 520, 683–7.

30. Mok, S., Ashley, E. A., Ferreira, P. E., Zhu, L., Lin, Z., Yeo, T., Chotivanich, K., Imwong, M., Pukrittayakamee, S. & Dhorda, M. (2015) Population transcriptomics of human malaria parasites reveals the mechanism of artemisinin resistance, Science. 347, 431–435.

31. Xue-Franzén, Y., Henriksson, J., Bürglin, T. R. & Wright, A. P. (2013) Distinct roles of the Gcn5 histone acetyltransferase revealed during transient stress-induced reprogramming of the genome, BMC genomics. 14, 1471–2164.

32. Gaupel, A. C., Begley, T. J. & Tenniswood, M. (2015) Gcn5 Modulates the Cellular Response to Oxidative Stress and Histone Deacetylase Inhibition, J Cell Biochem. 116, 1982–92.

33. Hu, Z., Song, N., Zheng, M., Liu, X., Liu, Z., Xing, J., Ma, J., Guo, W., Yao, Y., Peng, H., Xin, M., Zhou, D. X., Ni, Z. & Sun, Q. (2015) Histone acetyltransferase GCN5 is essential for heat stress-responsive gene activation and thermotolerance in Arabidopsis, Plant J. 84, 1178–91.

34. Johnsson, A., Xue-Franzen, Y., Lundin, M. & Wright, A. P. (2006) Stress-specific role of fission yeast Gcn5 histone acetyltransferase in programming a subset of stress response genes, Eukaryot Cell. 5, 1337–46.

35. Naguleswaran, A., Elias, E. V., McClintick, J., Edenberg, H. J. & Sullivan, W. J., Jr. (2010) Toxoplasma gondii lysine acetyltransferase GCN5-A functions in the cellular response to alkaline stress and expression of cyst genes, PLoS Pathog. 6, e1001232.

36. Fan, Q., An, L. & Cui, L. (2004) Plasmodium falciparum histone acetyltransferase, a yeast GCN5 homologue involved in chromatin remodeling, Eukaryot Cell. 3, 264–76.

37. Fan, Q., An, L. & Cui, L. (2004) PfADA2, a Plasmodium falciparum homologue of the transcriptional coactivator ADA2 and its in vivo association with the histone acetyltransferase PfGCN5, Gene. 336, 251–61.

38. Cui, L., Miao, J., Furuya, T., Li, X., Su, X. Z. & Cui, L. (2007) PfGCN5-mediated histone H3 acetylation plays a key role in gene expression in Plasmodium falciparum, Eukaryot Cell. 6, 1219–27.

39. Bartfai, R., Hoeijmakers, W. A., Salcedo-Amaya, A. M., Smits, A. H., Janssen-Megens, E., Kaan, A., Treeck, M., Gilberger, T. W., Francoijs, K. J. & Stunnenberg, H. G. (2010) H2A.Z demarcates intergenic regions of the plasmodium falciparum epigenome that are dynamically marked by H3K9ac and H3K4me3, PLoS Pathog. 6, e1001223.

40. Kanyal, A., Rawat, M., Gurung, P., Choubey, D., Anamika, K. & Karmodiya, K. (2018) Genome-wide survey and phylogenetic analysis of histone acetyltransferases and histone deacetylases of Plasmodium falciparum, FEBS J. 285, 1767–1782.

41. Zininga, T., Achilonu, I., Hoppe, H., Prinsloo, E., Dirr, H. W. & Shonhai, A. (2015) Overexpression, Purification and Characterisation of the Plasmodium falciparum Hsp70-z (PfHsp70-z) Protein, PLoS One. 10, e0129445.

42. Percario, S., Moreira, D. R., Gomes, B. A., Ferreira, M. E., Goncalves, A. C., Laurindo, P. S., Vilhena, T. C., Dolabela, M. F. & Green, M. D. (2012) Oxidative stress in malaria, International journal of molecular sciences. 13, 16346–72.

43. Bushell, E., Gomes, A. R., Sanderson, T., Anar, B., Girling, G., Herd, C., Metcalf, T., Modrzynska, K., Schwach, F., Martin, R. E., Mather, M. W., McFadden, G. I., Parts, L., Rutledge, G. G., Vaidya, A. B., Wengelnik, K., Rayner, J. C. & Billker, O. (2017) Functional Profiling of a Plasmodium Genome Reveals an Abundance of Essential Genes, Cell. 170, 260–272 e8.

44. Zhang, M., Wang, C., Otto, T. D., Oberstaller, J., Liao, X., Adapa, S. R., Udenze, K., Bronner, I. F., Casandra, D., Mayho, M., Brown, J., Li, S., Swanson, J., Rayner, J. C., Jiang, R. H. Y. & Adams, J. H. (2018) Uncovering the essential genes of the human malaria parasite Plasmodium falciparum by saturation mutagenesis, Science. 360.

45. Bridgford, J. L., Xie, S. C., Cobbold, S. A., Pasaje, C. F. A., Herrmann, S., Yang, T., Gillett, D. L., Dick, L. R., Ralph, S. A., Dogovski, C., Spillman, N. J. & Tilley, L. (2018) Artemisinin kills malaria parasites by damaging proteins and inhibiting the proteasome, Nat Commun. 9, 3801.

46. Zhang, M., Gallego-Delgado, J., Fernandez-Arias, C., Waters, N. C., Rodriguez, A., Tsuji, M., Wek, R. C., Nussenzweig, V. & Sullivan, W. J., Jr. (2017) Inhibiting the Plasmodium eIF2alpha Kinase PK4 Prevents Artemisinin-Induced Latency, Cell Host Microbe. 22, 766–776 e4.

47. Jeffers, V., Gao, H., Checkley, L. A., Liu, Y., Ferdig, M. T. & Sullivan, W. J. (2016) Garcinol inhibits GCN5-mediated lysine acetyltransferase activity and prevents replication of the parasite Toxoplasma gondii, Antimicrobial agents and chemotherapy. 60, 2164–2170.

48. Bhattacharjee, S., Coppens, I., Mbengue, A., Suresh, N., Ghorbal, M., Slouka, Z., Safeukui, I., Tang, H. Y., Speicher, D. W., Stahelin, R. V., Mohandas, N. & Haldar, K. (2018) Remodeling of the malaria parasite and host human red cell by vesicle amplification that induces artemisinin resistance, Blood. 131, 1234–1247.

49. Goyal, M., Alam, A., Iqbal, M. S., Dey, S., Bindu, S., Pal, C., Banerjee, A., Chakrabarti, S. & Bandyopadhyay, U. (2012) Identification and molecular characterization of an Alba-family protein from human malaria parasite Plasmodium falciparum, Nucleic acids research. 40, 1174–90.

50. Vembar, S. S., Macpherson, C. R., Sismeiro, O., Coppee, J. Y. & Scherf, A. (2015) The PfAlba1 RNA-binding protein is an important regulator of translational timing in Plasmodium falciparum blood stages, Genome Biol. 16, 212.

51. Verma, J. K., Wardhan, V., Singh, D., Chakraborty, S. & Chakraborty, N. (2018) Genome-Wide Identification of the Alba Gene Family in Plants and Stress-Responsive Expression of the Rice Alba Genes, Genes. 9.

52. Alves, L. R. & Goldenberg, S. (2016) RNA-binding proteins related to stress response and differentiation in protozoa, World journal of biological chemistry. 7, 78–87.

53. Karmodiya, K., Krebs, A.R., Oulad-Abdelghani, M., Kimura, H., Tora, L. (2012) H3K9 and H3K14 acetylation co-occur at many gene regulatory elements, while H3K14ac marks a subset of inactive inducible promoters in mouse embryonic stem cells, BMC genomics. 13, 1471–2164.

54. Kuo, M. H., vom Baur,E., Struhl, K., Allis, C.D. (2000) Gcn4 Activator Targets Gcn5 Histone Acetyltransferase to Specific Promoters Independently of Transcription, Molecular Cell. 6, 1309–1320.

55. Zhang, M., Mishra, S., Sakthivel, R., Rojas, M., Ranjan, R., Sullivan, W. J., Jr., Fontoura, B. M., Menard, R., Dever, T. E. & Nussenzweig, V. (2012) PK4, a eukaryotic initiation factor 2alpha(eIF2alpha) kinase, is essential for the development of the erythrocytic cycle of Plasmodium, Proc Natl Acad Sci U S A. 109, 3956–61.

56. Rao, R., Nalluri, S., Kolhe, R., Yang, Y., Fiskus, W., Chen, J., Ha, K., Buckley, K. M., Balusu, R. & Coothankandaswamy, V. (2010) Treatment with panobinostat induces glucose-regulated protein 78 acetylation and endoplasmic reticulum stress in breast cancer cells, Molecular cancer therapeutics, 1535–7163. MCT-09-0988.

57. Scherf, A., Hernandez-Rivas, R., Buffet, P., Bottius, E., Benatar, C., Pouvelle, B., Gysin, J. & Lanzer, M. (1998) Antigenic variation in malaria: in situ switching, relaxed and mutually exclusive transcription of var genes during intra-erythrocytic development in Plasmodium falciparum, The EMBO journal. 17, 5418–5426.

58. Deitsch, K. W. & Dzikowski, R. (2017) Variant gene expression and antigenic variation by malaria parasites, Annual review of microbiology. 71, 625–641.

59. Tonkin, C. J., Carret, C. K., Duraisingh, M. T., Voss, T. S., Ralph, S. A., Hommel, M., Duffy, M. F., da Silva, L. M., Scherf, A. & Ivens, A. (2009) Sir2 paralogues cooperate to regulate virulence genes and antigenic variation in Plasmodium falciparum, PLoS biology. 7, e1000084.

60. Coleman, B. I., Skillman, K. M., Jiang, R. H., Childs, L. M., Altenhofen, L. M., Ganter, M., Leung, Y., Goldowitz, I., Kafsack, B. F. & Marti, M. (2014) A Plasmodium falciparum histone deacetylase regulates antigenic variation and gametocyte conversion, Cell host & microbe. 16, 177–186.

61. Volz, J. C., Bártfai, R., Petter, M., Langer, C., Josling, G. A., Tsuboi, T., Schwach, F., Baum, J., Rayner, J. C. & Stunnenberg, H. G. (2012) PfSET10, a Plasmodium falciparum methyltransferase, maintains the active var gene in a poised state during parasite division, Cell host & microbe. 11, 7–18.

62. Jiang, L., Mu, J., Zhang, Q., Ni, T., Srinivasan, P., Rayavara, K., Yang, W., Turner, L., Lavstsen, T. & Theander, T. G. (2013) PfSETvs methylation of histone H3K36 represses virulence genes in Plasmodium falciparum, Nature. 499, 223.

63. Perez-Toledo, K., Rojas-Meza, A. P., Mancio-Silva, L., Hernandez-Cuevas, N. A., Delgadillo, D. M., Vargas, M., Martinez-Calvillo, S., Scherf, A. & Hernandez-Rivas, R. (2009) Plasmodium falciparum heterochromatin protein 1 binds to tri-methylated histone 3 lysine 9 and is linked to mutually exclusive expression of var genes, Nucleic acids research. 37, 2596–606.

64. Flueck, C., Bartfai, R., Volz, J., Niederwieser, I., Salcedo-Amaya, A. M., Alako, B. T., Ehlgen, F., Ralph, S. A., Cowman, A. F., Bozdech, Z., Stunnenberg, H. G. & Voss, T. S. (2009) Plasmodium falciparum heterochromatin protein 1 marks genomic loci linked to phenotypic variation of exported virulence factors, PLoS Pathog. 5, e1000569.

65. Chene, A., Vembar, S. S., Riviere, L., Lopez-Rubio, J. J., Claes, A., Siegel, T. N., Sakamoto, H., Scheidig-Benatar, C., Hernandez-Rivas, R. & Scherf, A. (2012) PfAlbas constitute a new eukaryotic DNA/RNA-binding protein family in malaria parasites, Nucleic Acids Res. 40, 3066–77.

66. Schroeder, M., Brooks, B. D. & Brooks, A. E. (2017) The Complex Relationship between Virulence and Antibiotic Resistance, Genes (Basel). 8.

67. Geisinger, E. & Isberg, R. R. (2017) Interplay Between Antibiotic Resistance and Virulence During Disease Promoted by Multidrug-Resistant Bacteria, J Infect Dis. 215, S9–S17.

68. Geisinger, E., Mortman, N. J., Vargas-Cuebas, G., Tai, A. K. & Isberg, R. R. (2018) A global regulatory system links virulence and antibiotic resistance to envelope homeostasis in Acinetobacter baumannii, PLoS Pathog. 14, e1007030.

69. Schneider, P., Chan, B. H., Reece, S. E. & Read, A. F. (2008) Does the drug sensitivity of malaria parasites depend on their virulence?, Malar J. 7, 257.

70. Radfar, A., Mendez, D., Moneriz, C., Linares, M., Marin-Garcia, P., Puyet, A., Diez, A. & Bautista, J. M. (2009) Synchronous culture of Plasmodium falciparum at high parasitemia levels, Nat Protoc. 4, 1899–915.

71. Zhang, Y., Liu, T., Meyer, C. A., Eeckhoute, J., Johnson, D. S., Bernstein, B. E., Nusbaum, C., Myers, R. M., Brown, M., Li, W. & Liu, X. S. (2008) Model-based analysis of ChIP-Seq (MACS), Genome Biol. 9, R137.

72. Quinlan, A. R. & Hall, I. M. (2010) BEDTools: a flexible suite of utilities for comparing genomic features, Bioinformatics. 26, 841–2.

73. Ye T, K. A., Choukrallah MA, Keime C, Plewniak F, Davidson I and Tora, L. (2011) seqMINER: an integrated ChIP-seq data interpretation platform, Nucleic Acids Res. 39, e35.

74. Witkowski, B., Amaratunga, C., Khim, N., Sreng, S., Chim, P., Kim, S., Lim, P., Mao, S., Sopha, C., Sam, B., Anderson, J. M., Duong, S., Chuor, C. M., Taylor, W. R. J., Suon, S., Mercereau-Puijalon, O., Fairhurst, R. M. & Menard, D. (2013) Novel phenotypic assays for the detection of artemisinin-resistant Plasmodium falciparum malaria in Cambodia: in-vitro and ex-vivo drug-response studies, The Lancet Infectious Diseases. 13, 1043–1049.

